# Chaperone-Mediated Autophagy in Fish: A Key Function Amid a Changing Environment

**DOI:** 10.1101/2024.03.20.585855

**Authors:** Simon Schnebert, Emilio J Vélez, Maxime Goguet, Karine Dias, Vincent Véron, Isabel García-Pérez, Lisa M Radler, Emilie Cardona, Stéphanie Fontagné-Dicharry, Pierre Van Delft, Franziska Dittrich-Domergue, Amélie Bernard, Florian Beaumatin, Amaury Herpin, Beth Cleveland, Iban Seiliez

## Abstract

Chaperone-Mediated Autophagy (CMA) is a major pathway of lysosomal proteolysis critical for cellular homeostasis and metabolism. While extensively studied in mammals, CMA’s existence in fish has only been confirmed recently, offering exciting insights into its role in species facing environmental stress. Here, we shed light on the existence of 2 genes encoding the CMA-limiting factor Lamp2A (lysosomal associated membrane protein 2A) in rainbow trout (RT, *Oncorhynchus mykiss*), revealing distinct expression patterns across various tissues. Notably, RT lacking the most expressed Lamp2A exhibit profound hepatic proteome disturbances during acute nutritional stress, underscoring its pivotal role as a guardian of hepatic proteostasis. Building upon these findings, we introduce and validate the CMA activation score as a reliable indicator of CMA status, providing a valuable tool for detecting cellular stress in fish under environmental threats. Overall, our study offers new perspectives into understanding CMA from evolutionary and environmental contexts.

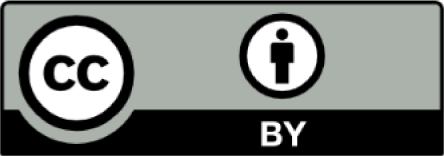

## INTRODUCTION

The higher frequency of extreme climate events is likely to have an impact on fish populations, production and fisheries by causing a shift in their physiology thus survival^1,2^. Parameters such as growth^3^, osmotic balance^4^, reproduction^5^, and cellular processes^6^ are directly affected. Cellular homeostasis is hence key for resisting environmental stressors coming fish’s way, and finding new ways of gauging it have become urgent.

In mammals, a lysosomal degradative process called autophagy involved in the recycling of proteins, damaged organelles and defense against pathogens is crucial to cellular homeostasis^7^. One of the selective forms of autophagy, chaperone-mediated autophagy (CMA), begins when cytosolic proteins bearing a KFERQ-like motif are recognized by the heat-shock protein family A HSP70 member 8 (HSPA8/HSC70) and brought to lysosomal membrane via specific binding to lysosome-associated membrane protein 2A (LAMP2A), the rate-limiting factor in CMA^8^. LAMP2A, one of the three splice variants (LAMP2A, B and C) of the *lamp2* gene, is the only one involved in CMA^9,10,11^. Following the binding of the substrate/chaperone complex, LAMP2A multimerizes to form a translocation complex through which the substrate translocates for intralysosomal degradation^12^. KFERQ-like motifs can be found in a large subset of the eukaryotic proteome in 2 different forms: canonical and putative. Putative motifs can be turned into KFERQ-like motifs after post-translational modifications such as phosphorylation or acetylation^13^. Proteins bearing canonical or putative KFERQ-like motifs have been associated with various biological processes like metabolism, cell cycle, immune response and transcriptional regulation^12^. Upon oxidative stress, nutrient deprivation, hypoxia or pathologies, enhanced CMA activity has been shown to regulate energetics and metabolic processes to maintain cellular homeostasis^14,15^. Fine tuning of the proteome and preservation of cellular homeostasis define CMA as a major cellular function in vertebrates.

Although previously thought to be restricted to mammals and birds, CMA was recently established in fish^16,17^. This discovery brought new perspectives for a better understanding of the mechanisms involved in the adaptation of organisms amid environmental stress. The rainbow trout (RT, *Oncorhynchus mykiss*), a widely spread salmonid commonly found in aquaculture production^18^, is an extensively studied model organism in physiology, genetics, behavior and ecology^19^. In addition to stressful environmental events it has to endure^20^, the RT also exhibits distinctive metabolic features like a high dietary protein requirement and glucose intolerance^21^. Therefore, it is an interesting candidate organism to study CMA and its potential role in regulating the proteome, particularly proteins responsible for maintaining metabolic and cellular homeostasis.

Recently, a homology-based search unveiled the presence of two *lamp2* paralogous genes within the genome of RT, located on chromosomes 14 (C14) and 31 (C31) respectively, with both genes harboring all three alternative exons (A, B, and C)^16,17^. Furthermore, a first glimpse of *lamp2a* splice variants expression in different fish species provided a clue to the potential existence of CMA in the RT^22^.

In this study, we have shown that RT has all the genetic material to express CMA machinery, particularly in tissues involved in major metabolic processes. We conducted a lysosome isolation combined with the Glyceraldehyde-3-phosphate Dehydrogenase (GAPDH) binding/uptake assay to measure CMA activity. It revealed that RT lysosomes can translocate GAPDH, a *bona fide* CMA substrate, depending on the presence of HSC70 and ATP. In addition, we showed that as in mammals, nutritional status regulates CMA activity. To fully unravel the physiological functions of CMA in RT, we used the genome-editing tool CRISPR-Cas9 and generated a knock-out (KO) line lacking LAMP2A from C31, the most expressed of both isoforms. Following LAMP2A C31 deletion, RTs were significantly larger than their wild type (WT) correlates and presented heavier livers and viscera weights. To further characterize the physiological roles of CMA in RT, we then used quantitative proteomics and found that upon nutritional stress, CMA is crucial to the preservation of cellular homeostasis by regulating oxidative stress response, energetics and metabolic processes. Finally, the adaptation and validation in RT of the “CMA score”, a reliable indicator of the activation status of CMA based on the expression levels of key CMA effectors and regulators as developed by Bourdenx et al. 2021^15^, not only introduces a novel biomarker for fish cellular homeostasis but also enables the assessment of CMA potential in emerging models. Overall, this study emphasizes the importance of considering CMA as a major function in maintaining fish cellular homeostasis, particularly amid the rising occurrence of new environmental threats.

## RESULTS

### The main CMA effectors are expressed in RT

A homology-based investigation of sequences of main CMA effectors in RT’s genome allowed us to investigate their expression throughout development and in different tissues of adult individuals. As presented in Fig 1A, most of core CMA genes are gradually induced from oocyte stage to maximum expression at stage 22/23, during which the primitive liver and the primitive hepatic portal vein develop in RT^23^. Both *lamp2a* paralogs from C14 and C31 were then compared and in most tissues, *lamp2a* C31 shows a higher expression (Fig 1B). Nearly all tissues displayed *lamp2a* expression, particularly liver, intestine and kidney.

**Figure 1.**
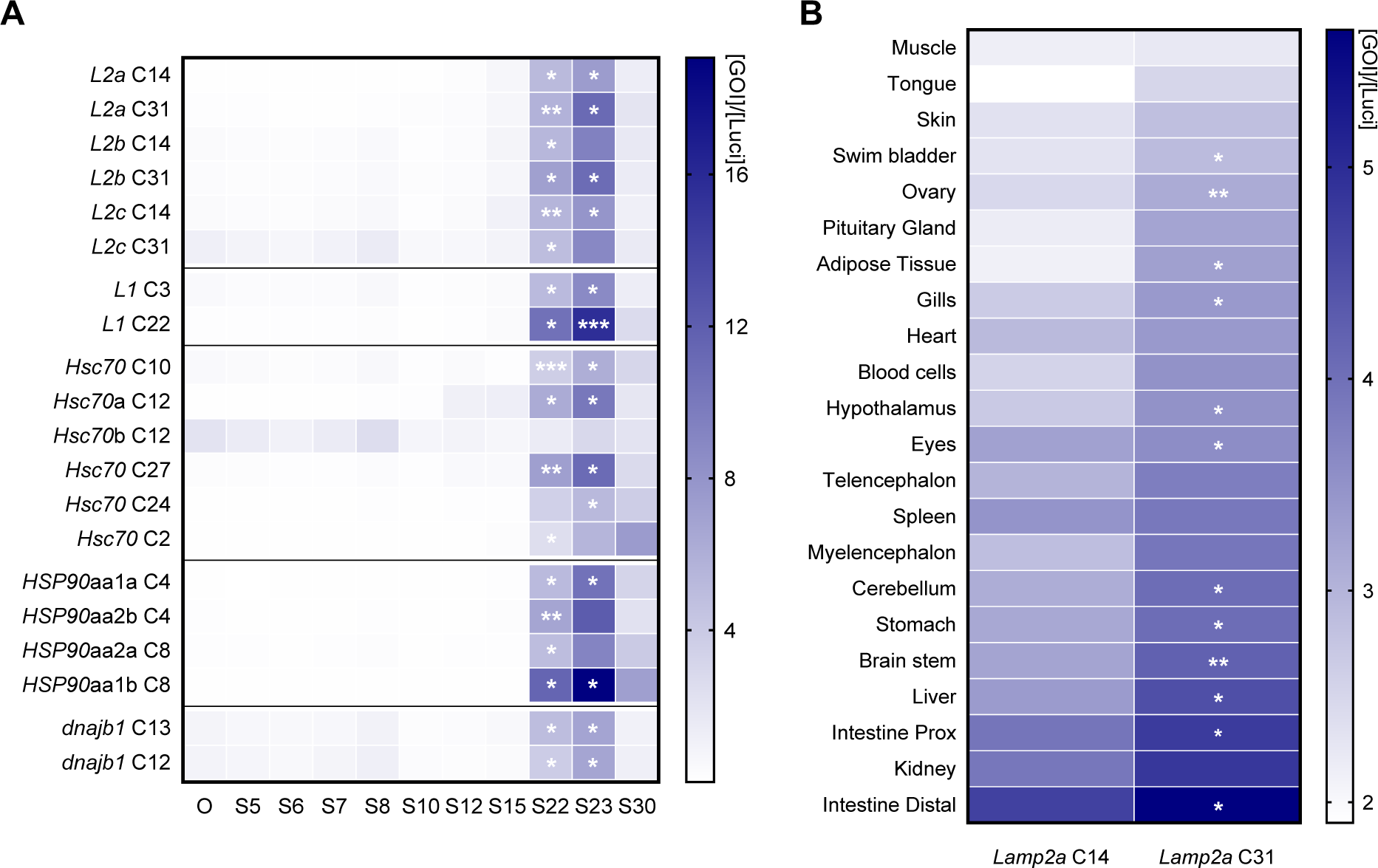
The expression of CMA-related genes in RT is induced during development and shows a tissue-specific pattern. (A) Expression of CMA-related genes throughout development from oocyte (O) stage (S) to hatching (S30). mRNA levels were normalized with luciferase. Parametric statistical t-tests were performed comparing all stages to oocyte stage (n=3 pools of embryos, one pool of 30 embryos per spawn) (*, *p*<0.05; **, *p*<0.01; ***, *p*<0.001). (B) Expression of *lamp2a* paralogs (from C14 and C31) in different tissues. mRNA levels were normalized with luciferase. A parametric statistical t-test was performed comparing both *lamp2a* paralogs (n=6) (*, *p*<0.05; **, *p*<0.01).

Following the high mRNA expression of both *lamp2a* paralogs in liver and intestine, we conducted a BaseScope assay to address spatial expression of the transcripts. *Lamp2a* transcripts quantity and distribution were analyzed in fed and starved RT, as starvation induces maximal CMA activation^24^. BaseScope signal showed that mRNA was distributed evenly across sampled liver tissue (Fig 2A). Countings confirmed previous data validating a higher expression of *lamp2a* C31 compared to *lamp2a* C14, regardless of nutritional status (Fig 2C).

**Figure 2.**
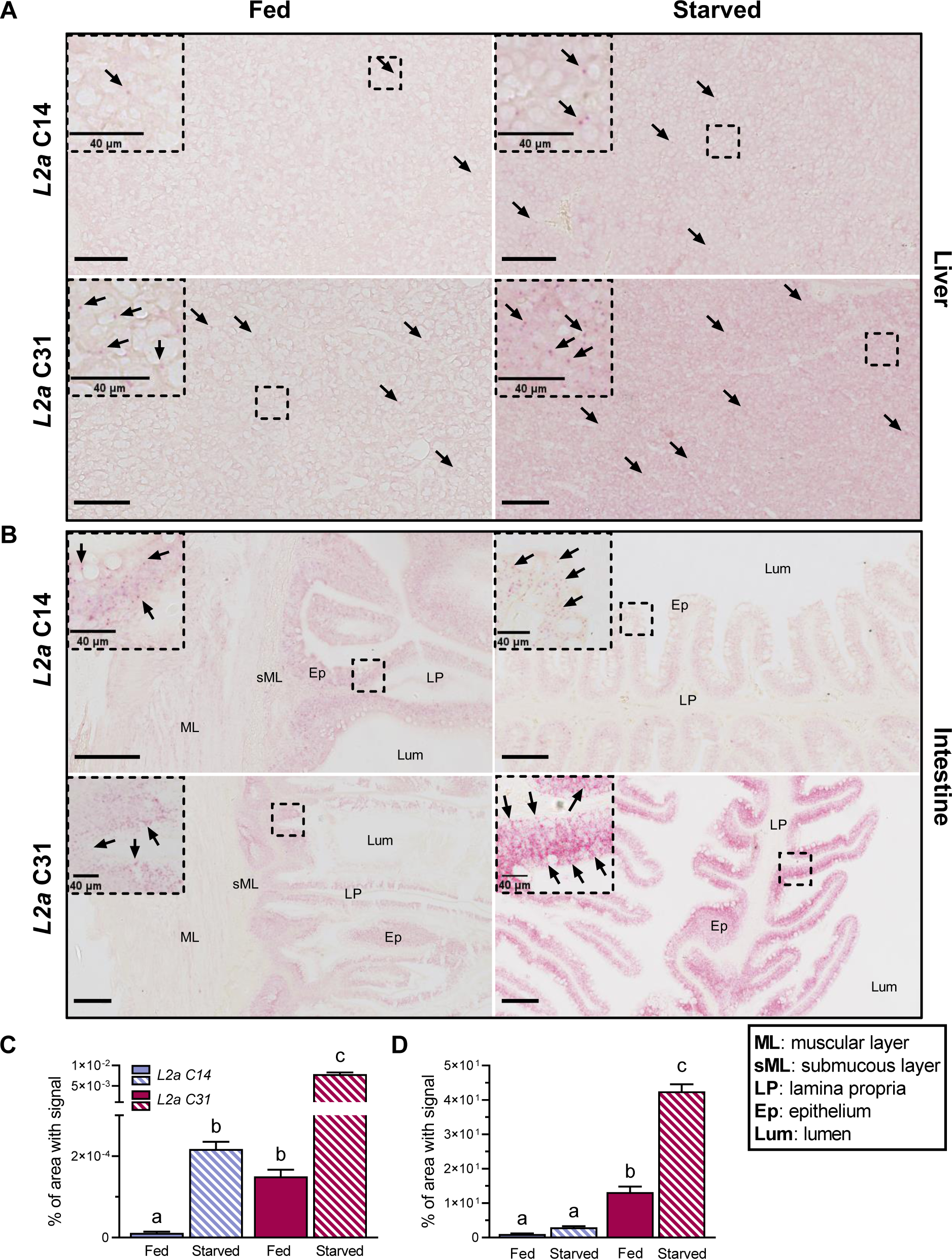
The expression of both RT lamp2a is induced by fasting and shows a similar spatial distribution although at different levels. (A) Signal of *lamp2a* C14 and *lamp2a* C31 probes (arrows) in liver sections from fed and starved RTs (Scale bars, 200 μm). (B) Signal of *lamp2a* C14 and *lamp2a* C31 probes (arrows) in distal intestine sections from fed and starved RTs (Scale bars, 200 μm). Enlargements of the boxed region are shown to the top left of each image (Scale bar, 40 µm). (C) Quantification of liver mRNA signal between both paralogs in contrasted nutritional states (n=4). (D) Quantification of distal intestine mRNA signal between both paralogs in contrasted nutritional states (n=4). A Kruskal-Wallis test (C) and a one-way ANOVA (D) were conducted comparing all conditions. Letters indicate a significant difference between treatments (*p*<0.001). All data is presented as mean +/- SEM.

In the distal part of the intestine, *lamp2a* mRNA was mostly found in enterocytes (epithelium) in contrast to lamina propria, serous and submucous layer (Fig 2B). Regarding the expression of *lamp2a* paralogs and effect of nutritional status, similar results were found in the intestine, with a higher expression of both *lamp2a* paralogs in starved conditions and *lamp2a* C31 being the most expressed transcript (Fig 2D).

The expression of CMA-related genes showed undisputable signs that RT carries all the genetic material required to express the CMA machinery, although a demonstration of its activity was required.

### RT exhibits CMA-active lysosomes able to bind and uptake GAPDH

To truly grasp whether CMA is functional in RT, we referred to the gold standard for *in vivo* CMA activity quantification, which consists of measuring the binding and uptake of a given CMA substrate in isolated CMA-competent (CMA+) lysosomes^25^. In this aim, we relied on the technique described by Juste & Cuervo’s^26^ to isolate trout liver lysosomes. Four interphases were separated after the ultra-centrifugation step theoretically containing, from top to bottom: CMA+ lysosomes, a mix population of CMA-competent and CMA-non-competent lysosomes (CMA+/- lysosomes), a mixture of lysosomes and mitochondria (Lyso/Mito) and mitochondria (Mito) (Fig 3A).

**Figure 3.**
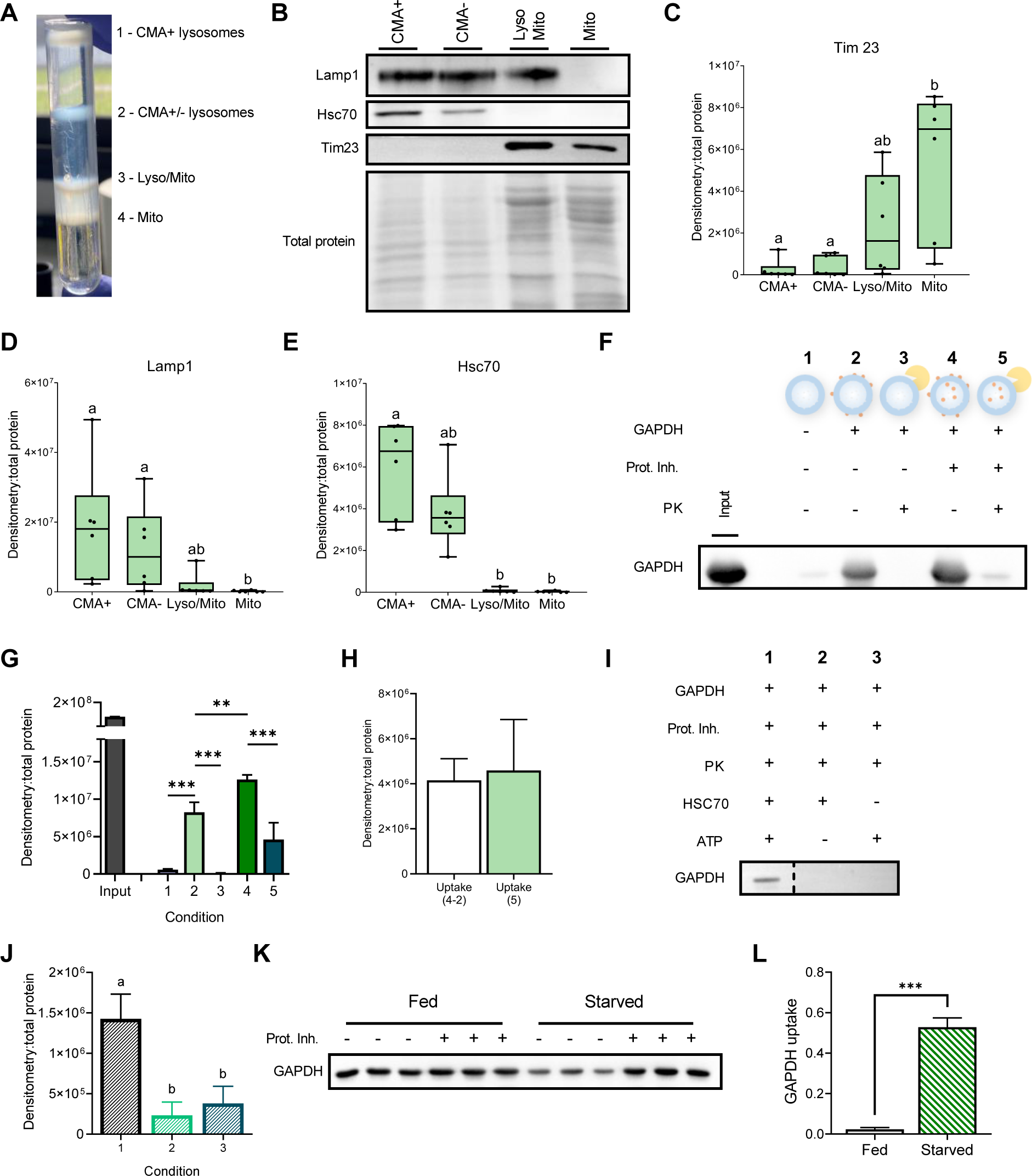
RT displays CMA-active lysosomes able to bind and uptake the CMA substrate GAPDH. (A) Different interphases obtained after centrifugation of RT liver homogenate on a discontinuous Nycodenz gradient. The mitochondria (Mito) are localized in the lowest interphase of the gradient, while the subsequent interphase consists of a mixture of lysosomes and mitochondria (Lyso/Mito). The third interphase from the bottom contains a heterogeneous population of CMA-competent and CMA-non-competent lysosomes (CMA+/- lysosomes), and the lysosomes with high CMA activity (CMA+) migrate to the upper interphase. (B) Immunoblots in those four different liver fractions and densitometric quantification of TIM23 (C), LAMP1 (D), and HSC70 (E) proteins. All values are shown as min. to max. (n=6). A Friedman test with multiple comparisons was performed comparing all fractions. Letters indicate a significant difference between treatments (*p*<0.0001). (F) Immunoblot showing RT CMA+ lysosome’s ability to bind and uptake GAPDH. Conditions correspond to (1) untreated CMA+ lysosomes, (2) CMA+ lysosomes incubated with the CMA substrate GAPDH, (3) CMA+ lysosomes incubated with GAPDH and PK, (4) CMA+ lysosomes incubated with GAPDH and inhibitors of lysosome proteases, and (5) CMA+ lysosomes incubated with GAPDH, Prot. Inh. and PK. (G) Densitometric quantification of the Western blot shown in F. Values are expressed relative to total proteins (n=6). A t-test (**, *p<*0.01; ***, *p<*0.001) was conducted to stress the differences between association, binding and uptake. (H) Densitometric quantification of GAPDH uptake, comparing uptake from condition 5 and the uptake obtained from the arithmetic difference between condition 2 and 4. (I) Western blot showing RT’s CMA activity depends on presence of Hsc-70 and ATP. (J) Densitometric quantification of GAPDH uptake in absence of Hsc70 and ATP, statistically compared with a one-way ANOVA (*p<*0.01**), (n=4). (K and L) GAPDH binding and uptake upon nutrient deprivation versus control condition. Representative Immunoblot (K) and quantification of GAPDH uptake (L) calculated as the difference between GAPDH association (+ Prot. inh.) and GAPDH binding (-Prot. inh.). A t-test was conducted comparing both nutritional conditions (*p<*0.001***), (n=3). All data is presented as mean + SEM.

To ensure that each interphase contained its expected content, we conducted western blotting to quantify key proteins from each fraction. We first quantified Tim23, a component of the mitochondrial inner membrane^27^. The mitochondrial fraction displayed a higher level of Tim23 compared to other phases (Figs 3B and C). We then quantified levels of lysosome-associated membrane protein 1 (Lamp1), commonly used as a lysosomal marker^28^ as well as the CMA-key Hsc70 chaperone. We found a higher abundance of Lamp1 in fractions that contained lysosomes (Figs 3B and 3D). Interestingly, Hsc70 was present at higher levels in the CMA+ fraction than in its CMA-counterpart (Figs 3B and 3E), in accordance with the higher content of lysosomal Hsc70 in CMA+ lysosomes^28,29^. Overall, these results clearly show that RT contains lysosomes displaying characteristics of CMA+ lysosomes.

After confidently having isolated CMA+ lysosomes, we needed to find evidence for their capacity to perform CMA. To assess CMA activity, we referred to the Juste & Cuervo’s protocol, which involves testing the ability of a given CMA substrate (e.g., GAPDH) to bind to and be taken up by CMA+ lysosomes^26^. Briefly, 5 different conditions were used to test CMA activity in RT (Fig 3F; quantification in Fig 3G). In condition 1, CMA+ lysosomes were left untreated. The thin GAPDH band observed by western blotting likely stems from endogenous RT GAPDH. In condition 2, we incubated CMA+ lysosomes with GAPDH. This resulted in a more pronounced band, indicating the binding of the CMA substrate to the lysosomal membrane (Figs 3F and 3G). To confirm the binding of GAPDH to the lysosomal membrane, we then introduced proteinase K (PK) in condition 3, which degraded all proteins associated with the cytosolic side of the lysosomal membrane. Consequently, the band disappeared, validating that GAPDH binds specifically to the lysosomal membrane (Figs 3F and 3G). In condition 4, we pre-treated CMA+ lysosomes with inhibitors of lysosomal proteases to prevent the degradation of translocated substrates. This led to a stronger band compared to condition 2 (Figs 3F and 3G), providing evidence that GAPDH binds to the lysosomal membrane and is subsequently translocated inside CMA+ lysosomes. Lastly, in condition 5, CMA+ lysosomes were pre-treated with inhibitors of lysosomal proteases and then incubated with PK (Figs 3F and 3G). A thinner band appeared, which likely represents the fraction of GAPDH taken up by CMA+ lysosomes. Additionally, we were able to measure the uptake by calculating the difference in GAPDH quantity between conditions 4 (association) and 2 (binding) or through the use of PK (condition 5). No difference was found between both uptake values (Fig 3H), strengthening our results and supporting that RT exhibits CMA+ lysosomes able to bind and uptake GAPDH.

To characterize the mechanisms at play more comprehensively, we then verified whether GAPDH binding and uptake by CMA+ lysosomes depend on the presence of ATP and HSC70, as demonstrated in mammals^25^. We incubated freshly isolated RT liver CMA+ lysosomes with all factors required for binding/uptake of GAPDH and added PK to reveal uptake (Fig 3I, condition 1). In conditions 2 and 3, we repeated the treatments, but without ATP or HSC70, respectively. Our findings clearly demonstrate that GAPDH uptake was significantly diminished (if not completely suppressed) in the absence of ATP or HSC70 (Figs 3I and 3J), witnessing the essential part they play in the ability of RT CMA+ lysosomes to bind and translocate the added GAPDH.

Finally, since nutrient deprivation commonly induces CMA in mammals^24^ and higher *lamp2a* gene expression was found in starved RT (Figs 2A – 2D), we investigated the effects of starvation on GAPDH uptake by RT CMA+ lysosomes. Our findings revealed that CMA+ lysosomes isolated from starved RT exhibited a significantly higher ability to uptake GAPDH compared to lysosomes from fed RT (Figs 3K and 3L), hinting at a higher CMA activity following nutrient deprivation akin to that occurring in mammals.

Taken together, these results indicate that RT contains lysosomes able to perform CMA or CMA-like activity.

### Lamp2a C31 deletion leads to enhanced body size, altered organ indices, and metabolic dysregulation in RT

Having demonstrated the existence of CMA activity in RT, we then focused on unraveling its physiological role. Using the genetic tool CRISPR-Cas9, we generated RT models knocked-out for *lamp2a* C31 (hereafter referred to as L2AC31-KO), as it is the most expressed of both paralogs (Figs 1B and 2A - 2D), and *lamp2a* C14’s contribution to CMA appears marginal, as seen in our recent *in vitro* study^17^. More specifically, the generated fish line displayed a deletion of a 322 bp region in exon 9 of the *lamp2* gene (from C31) that encodes for the specific cytosolic and transmembrane domains of the Lamp2a protein (Fig 4A and S1A). As expected, levels of *lamp2a* C31 mRNA were undetectable in the tissues of L2AC31-KO RTs (Fig 4B). Surprisingly, higher expression of *lamp2a* C14 mRNA was found in L2AC31-KO RTs when compared to their WT counterparts (Fig 4C), suggesting the existence of compensatory mechanisms. Differential regulations of other *lamp2* splice variants transcripts (*lamp2b* and *lamp2c*) from both C31 and C14 were also observed (Figs S1B - S1E), in agreement with the reported specific regulation of expression of these 2 isoforms^30^, which do not participate in CMA.

**Figure 4.**
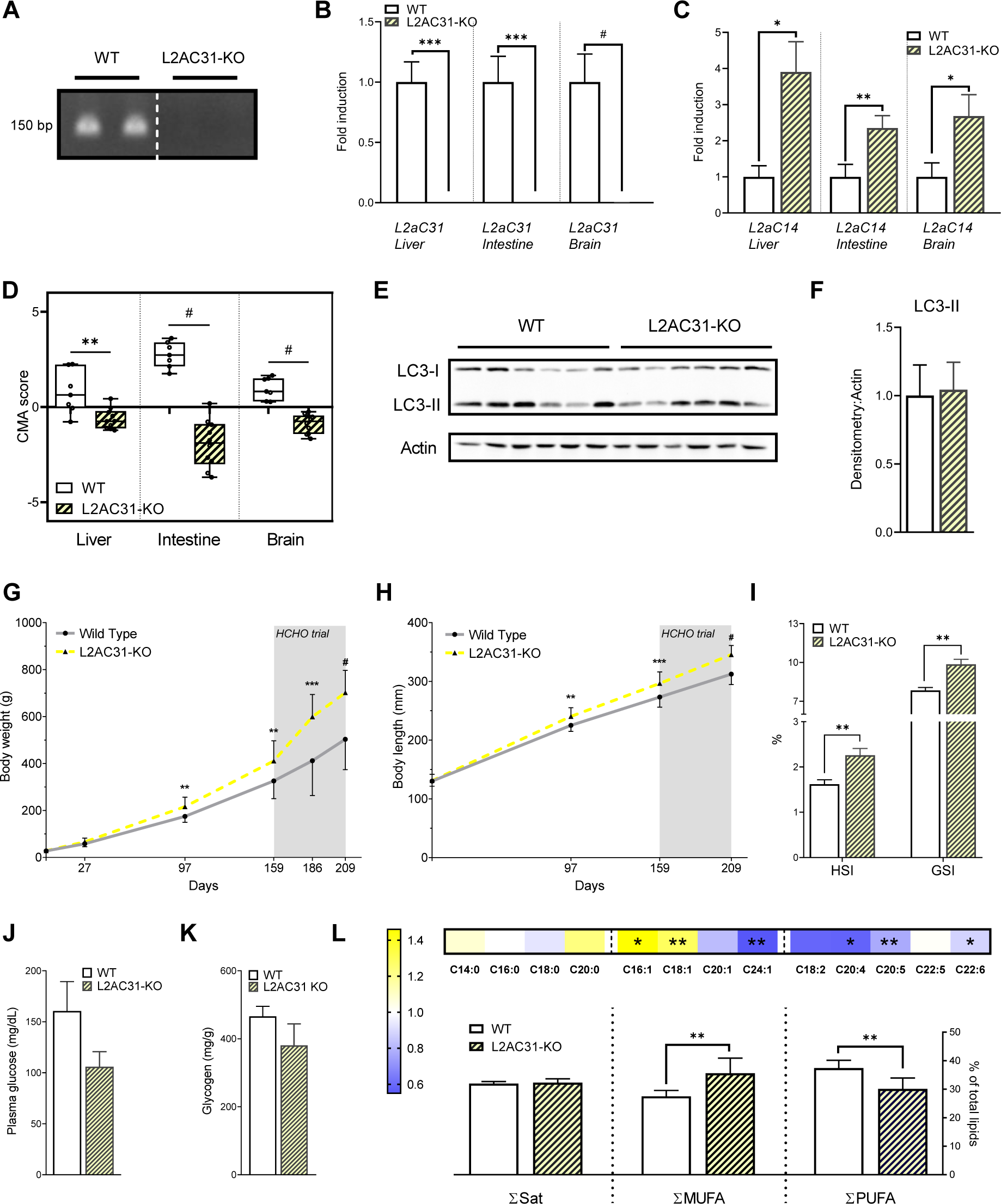
Lamp2a C31 deletion leads to enhanced body size, altered organ indices, and metabolic dysregulation in RT. (A) Genotyping of lamp2a allele was performed by PCR of fish fin generated by Crispr-Cas9 method. The primers used were specifically designed to bind exon A. (B) Liver, intestine and brain mRNA expression of *lamp2a* C31 in WT and L2AC31-KO RTs analyzed by RT-qPCR (n=7, values are mean + SEM shown by fold induction respect to WT fish). (C) Liver, intestine and brain mRNA expression of *lamp2a* C14 genes in WT vs L2AC31-KO RTs (n=7, values are mean + SEM shown by fold induction respect to WT fish). A Mann-Whitney test was performed comparing mRNA of WT and L2AC31-KO (*, *p<*0.05; **, *p<*0.01; ***, *p<*0.001; #, *p<*0.0001). (D) CMA score based on expression of liver, intestine and brain CMA network genes in WT vs L2AC31-KO RTs. (n=7, all values shown min. to max.) (t-test; **, *p<*0.01; #, *p<*0.0001). (E) Representative immunoblot for LC3 I and LC3 II in homogenate liver from WT and L2AC31-KO RTs, quantified in (F), values are mean + SEM. (G) Body weights of WT *vs* L2AC31-KO RTs during 209 days of experiment. Mean +/- SD (*, *p<*0.05; **, *p<*0.01; ***, *p<*0.001; #, *p<*0.0001). (H) Body lengths of L2AC31-KO vs WT RTs during 209 days of experiment. Mean +/- SD (*, *p<*0.05; **, *p<*0.01; ***, *p<*0.001; #, *p<*0.0001). (I) Hepatosomatic ([liver weight/body weight]x100) and Gastrosomatic index ([weight of gastrointestinal tract /body weight]x100) of WT *vs* L2AC31-KO RTS sampled at day 209. Values are mean + SEM (**, *p<*0.01). (J) Postprandial plasma glucose in WT *vs* L2AC31-KO RTs at day 209. Values are mean + SEM. (K) Hepatic glycogen content (mg of glycogen per g of fresh tissue) of WT *vs* L2AC31-KO RTs at day 209. Values are mean + SEM. (L) Hepatic lipid profile of WT and L2AC31-KO RTs at day 209. The heatmap represents the % of each fatty acid type to total lipids in L2AC31-KO divided by the corresponding value from the WT group. In both groups, a sum of percentages was calculated per fatty acid category: saturated (Sat), monounsaturated (MUFA) and polyunsaturated (PUFA). Values are mean + SEM. A t-test was performed comparing each fatty acid content between groups (**, *p<*0.01).

To further appreciate the impact of the *lamp2a* C31 deletion on the overall CMA network, we then used the CMA score, as recently defined by Bourdenx et al^15^. The score is a weighted average of the expression level of every known effector and both positive and negative modulators of CMA (Fig S2A and Table S1), with recent studies showing that it represents a reliable proxy for CMA activation status^29,31,32^. We first validated this score experimentally using RT cells expressing the KFERQ-PA-mCherry1 CMA reporter, a widely utilized method for monitoring CMA in mammalian cells^33^ and more recently validated in fish cells^16,17^. These cells were cultured under conditions known to induce CMA activation: starvation (serum-free) and mild-oxidative stress (H2O2, 25 µM) (Fig S2B). CMA activity (number of puncta/cell) was quantified after 4, 8, 16, 24 and 48h of treatment. The highest CMA activation was observed upon mild-oxidative stress and starvation at 16 hours (Figs S2C and S2D). The CMA score was then calculated at that time point using mRNA levels of all CMA network elements (Fig S2E). The significant higher CMA score encountered upon these 2 CMA activators respect to the control condition (Fig S2F) ensured its adequacy and the opportunity to use it as a potential proxy for CMA’s functional output in RT as well. After we validated the CMA activation score *in vitro*, we then calculated it for liver, intestine and brain samples of WT versus L2AC31-KO RTs. We found significant decrease of the CMA score in L2AC31-KO RTs in the different tissues analyzed (Fig 4D), suggesting a decline in CMA activity following *lamp2a* C31 KO. Interestingly, we found no significant alteration in the steady state levels of LC3-II, a well-established marker for autophagosome formation^34^, in the livers of L2AC31-KO RTs compared to WT fish (Figs 4E and 4F), highlighting the suitability of these animals to study the consequences of compromised CMA activity independent of any effects induced by macroautophagy compensation.

Considering the critical role(s) of CMA in the control of hepatic metabolism^35,16^, we then investigated the physiological consequences of *lamp2a* C31 deletion. In a first phase, we tracked the biometrical parameters/growth of both L2AC31-KO and WT fish for approximately 23 weeks, and observed that L2AC31-KO animals presented both significantly higher body weight and length than their WT counterparts both at 97 and 159 days of rearing (Figs 4G and 4H), suggesting that the lack of Lamp2a C31 induces metabolic alterations in these species. To go further, and as the RT is considered a natural model for glucose intolerance^36^, we then subjected both L2AC31-KO and WT fish to a diet rich in carbohydrates (HCHO diet, 30% of the diet). This dietary challenge aimed to investigate the nutritional stress response of RTs unable to perform CMA. Interestingly, after 7 weeks of feeding the high carbohydrate diet, L2AC31-KO fish still presented significantly higher body weights and lengths than WT (Figs 4G and 4H). Furthermore, at the last time point, L2AC31-KO fish exhibited significantly higher levels of hepatosomatic (HSI) and gastrosomatic (GSI) indexes (Fig 4I), as well as a trend towards reduced postprandial plasma glucose levels (Fig 4J), suggesting defects in their metabolism. Given the important phenotypical consequences observed in L2AC31-KO RTs, we inspected the specific metabolic pathways affected by *lamp2a* C31 inactivation. We found that glycogen levels were unaltered upon *lamp2a* C31 inactivation (Fig 4K), indicating that the liver enlargement in L2AC31-KO RTs was not caused by accumulated glycogen. Accordingly, we measured the hepatic lipid content and found similar levels of saturated fatty acids (Sat) in WT and L2AC31-KO RTs (Fig 4L). However, L2AC31-KO RTs displayed an accumulation of monounsaturated fatty acids (MUFA) and a depletion in polyunsaturated fatty acids (PUFA) (Fig 4L), a fatty acid composition found in nonalcoholic fatty liver^37^.

### Lamp2a C31 deletion in RT leads to substantial remodeling of the hepatic proteome

To unveil how Lamp2a C31 deficiency leads to the phenotypic changes observed, we first measured the expression of select genes involved in glucose and lipid metabolism. We found that mRNA expression levels of glucose metabolism-related enzymes in the livers of WT and L2AC31-KO RTs did not differ significantly (Fig 5A). In contrast, L2AC31-KO fish exhibited a significant upregulation of the genes involved in lipid metabolism (Fig 5A), hinting at an enhanced potential to synthesize fatty acids.

**Figure 5.**
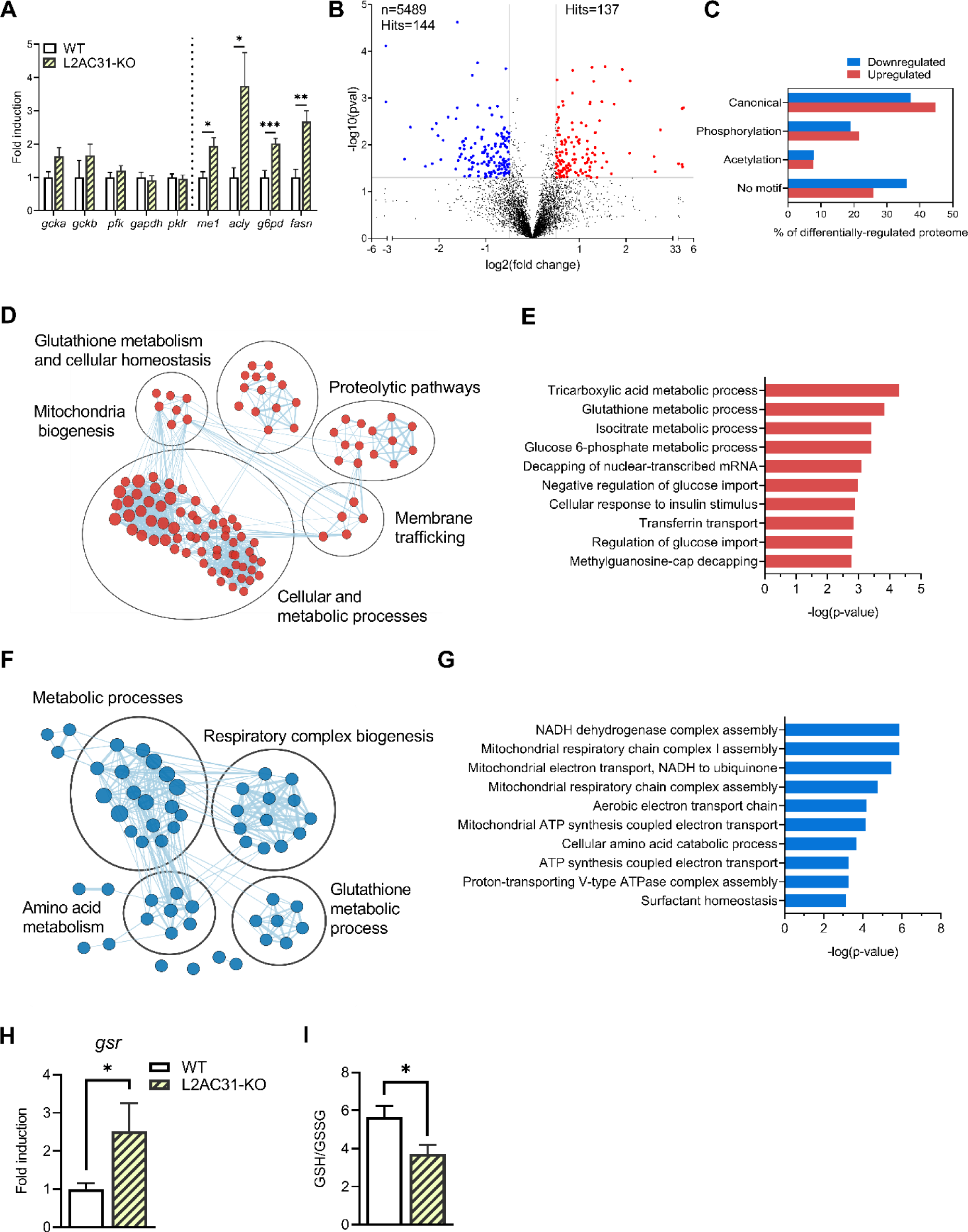
Deletion of Lamp2a C31 in RT leads to substantial remodeling of the hepatic proteome. (A) mRNA levels of several enzymes related to glucose and lipid metabolisms in liver homogenates from WT *vs* L2AC31-KO RTs (n=7, values are mean + SEM fold induction respect to WT group). *gck*a, *gck*b, glucokinase a and b; *pfk*, phosphofructokinase 1; *gapdh*, glyceraldehyde 3-phosphate dehydrogenase; *pklr*, pyruvate kinase; *me1*, malic enzyme 1; *acly*, ATP citrate lyase; *g6pd*, glucose-6-Phosphate dehydrogenase; *fasn*, fatty acid synthase. A Mann-Whitney test was performed comparing mRNA of WT and L2AC31-KO (*, *p<*0.05; **, *p<*0.01; ***, *p<*0.001). (B) Volcano plot of the quantitative proteomics analysis of liver L2AC31-KO VS WT RTS. Top left: number of identified proteins (n) and significantly down-regulated hits. Top right: number of significantly up-regulated hits. Blue and red dots indicate significantly down- and up-regulated proteins, respectively (*p<*0.05 and fold change >1.41). (C) Proportion of differentially-regulated proteins harboring distinct classes of KFERQ-like motifs. (D) Enrichment map showing biologically-related protein networks up-regulated upon Lamp2a blockage. (E) Gene ontology analysis of proteins up-regulated upon Lamp2a blockage. (F) Enrichment map showing biologically-related protein networks down-regulated upon Lamp2a blockage. (G) Gene ontology analysis of proteins down-regulated upon Lamp2a blockage. (H) Glutathione-Disulfide Reductase mRNA expression in liver L2AC31-KO *VS* WT RTS. Values are mean + SEM (n=7) shown by fold induction respect to WT group (Mann-Whitney test; *, *p<*0.05). (I) Hepatic GSSG:GSH level in L2AC31-KO VS WT RTS. Values are mean ± SEM (n=7; Mann-Whitney test; *, *p<*0.05).

To investigate whether L2AC31-KO RTs also underwent liver proteome perturbations, we conducted a quantitative proteomic analysis of the liver samples obtained from both WT and L2AC31-KO fish fed the HCHO diet. Overall, we identified 5489 proteins, among which 2.5% (137 proteins) were significantly up-regulated and 2.6% (144 proteins) were significantly down-regulated in L2AC31-KO RTs compared to WT (Fig 5B, Tables S2 and S3). Remarkably, the analysis of CMA-targeting motifs in these up- and down-regulated proteins revealed a higher percentage of proteins containing either the canonical or a phosphorylation-generated KFERQ motif in the former group compared to the latter group (Fig 5C), potentially suggesting an accumulation of putative CMA substrates in L2AC31-KO RTs.

To obtain a global picture of pathways affected by Lamp2a inactivation in RT, we first conducted an enrichment map analysis. We found that proteins up-regulated in L2AC31-KO RTs, were mainly related to cellular and metabolic processes (Fig 5D and Table S2). The gene ontology analysis then provided us with a closer look at specific pathways affected by Lamp2A C31 disruption (Fig 5E). Notably, among the proteins that were up-regulated, the biological processes most prominently observed were the TCA cycle and pathways associated with glucose and isocitrate metabolism. In detail, we found several proteins involved in the metabolic use of glucose as well as fatty acid biosynthesis and transport (Table S2). These include notably (i) hexokinase (HK) and fructose bisphosphate A (ALDOA) involved in glycolysis, (ii) isocitrate dehydrogenase (IDH) and malate dehydrogenase (MDH) involved in the TCA cycle, and (iii) ATP citrate synthase (ACLY), very long-chain acyl-CoA synthetase (ACSVL) and fatty-acid-binding proteins (FABP) involved in fatty acid biosynthesis and transport. Although this analysis cannot reveal whether these proteins (whose levels are induced in L2AC31-KO versus WT fish) are directly targeted by CMA, it is worth noting that a number of them have already been described as *bona fide* CMA substrates in mammals^38^ and possess one or more KFERQ motifs in RT (Table S4). Whether this upregulation of proteins involved in hepatic metabolism is direct or indirect, it supports an enhanced capacity of L2AC31-KO RT liver for glucose utilization and fatty acid biosynthesis, in agreement with the already mentioned reduced glucose circulating levels and altered hepatic lipid profile (Figs 4J and 4L). However, a significant number of proteins associated with metabolic processes were also found to be downregulated in KO fish compared to their WT counterparts (Fig 5F and Table S3). Among them, we noticed the presence of Apo lipoprotein B100 (ApoB-100), which is involved in the assembly and secretion of very low density lipoprotein (VLDL) associated with the distribution of excess triglycerides from the liver to peripheral tissues^39^. Thus, this finding supports defects in the hepatic lipid metabolism/trafficking.

Another cluster identified through enrichment mapping and gene ontology analysis is associated with mitochondria biogenesis and function (Figs 5D - 5G). Despite an increase in proteins related to mitochondrial biogenesis (Fig 5D and Table S2), we observed a decrease in the levels of electron transport chain complexes (Figs 5F and 5G, and Table S3). Specifically, various subunits of mitochondrial complex I (FOXRED1, NDUFV1, NDUFA7, NDUFA12) and complex IV (COX-7c) were found to be downregulated upon Lamp2a C31 inactivation (Table S3), suggesting potential alterations in mitochondrial respiration within the liver of L2AC31-KO RTs. Notably, our analysis also revealed elevated levels of several proteins associated with glutathione (GSH) metabolism and antioxidant functions in mutant fish, including glutamate cysteine ligase (GCL), Glutathione S-transferase omega (GSTO1), thioredoxin, thioredoxin reductase (TR), glutaredoxin, and glucose-6-phosphate dehydrogenase (G6PD) (Fig 5D and Table S2), possibly reflecting the establishment of an adaptive response to mitigate oxidative stress caused by a compromised mitochondrial function. In that direction, further investigation unveiled an upregulation of Glutathione-Disulfide Reductase (*gsr*) mRNA expression along with a decrease in the ratio of reduced glutathione (GSH) to oxidized glutathione (GSSG) in the liver of L2AC31-KO RTs (Figs 5H and 5I), supporting a situation of compromised redox status in CMA-defect animals.

Finally, we turned our attention to the cluster associated with proteolytic pathways evidenced by the enrichment map of the up-regulated proteins (Fig 5D). We witnessed an increase of key components of the endosomal sorting complexes required for transport (ESCRT) machinery such as vacuolar protein sorting 4 homolog B (VPS4B) (Table S2), considered to be an ESCRT accessory protein whose expression level controls the activation of endosomal microautophagy (eMI). Other proteins related to the ubiquitin-proteasome system (UPS), including Proteasome 26S subunit, ATPase 1 (PSMC1), UV excision repair protein RAD23 homolog A (RAD23) and Ubiquitin Conjugating Enzyme E2 K UBE2K, were also up-regulated in L2AC31-KO RTs (Table S2), overall suggesting that compensation of the CMA defect could occur via these two pathways.

Taken together, these results demonstrate that upon an acute nutritional stress such as the HCHO diet, loss of CMA leads to strong perturbations of the hepatic proteome, thus highlighting its critical role as a gatekeeper of hepatic proteostasis in RT.

### Use of CMA score as a prognostic tool to assess cellular homeostasis in fish exposed to hypoxic conditions

After emphasizing the critical role of CMA in the regulation of hepatic proteostasis in RT exposed to an acute stress, we investigated the applicability of the CMA score as a tool to assess cellular homeostasis in fish facing environmental stress. In this aim, we subjected RTs to moderate hypoxic conditions (dissolved oxygen DO 5.6 ± 0.5 ppm), widely known to induce oxidative stress and organ failure in fish, as established by previous studies^40,41^. After 4 weeks of moderate hypoxia, we collected the liver, gills and multiple brain parts of RT and calculated the CMA score (Fig 6A). We found a significant overall increase in the CMA score in RT tissues when exposed to hypoxia, suggesting the overall induction of the CMA network in response to this stressful condition (Fig 6B). Collectively, these findings highlight the activation of the CMA machinery following an environmental stress and emphasize the purpose of using the CMA score as an indicative marker for cellular homeostasis in organisms currently exposed to a changing environment.

**Figure 6.**
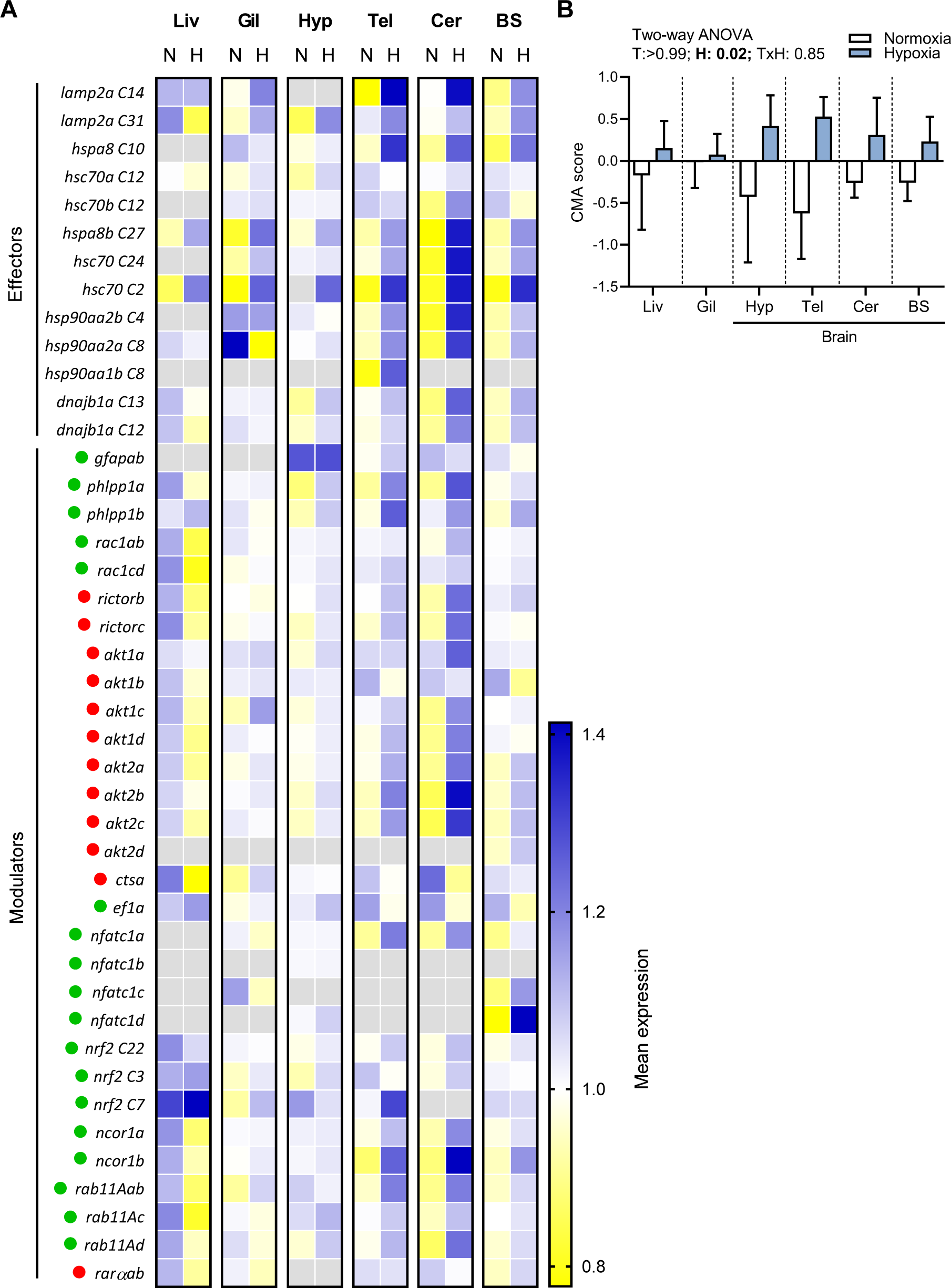
Use of CMA score as a prognostic tool to assess cellular homeostasis in fish exposed to stressful conditions. (A) Expression of CMA network genes in different tissues of RT under hypoxia relative to normoxia (n=6). N= normoxia; H= hypoxia; Liv= liver; Gil= gills; Hyp= hypothalamus; Tel= telencephalon; Cer= cerebellum; BS= brain stem. The CMA network components are organized into functional groups. Green and red dots indicate whether the element has a positive or negative impact on CMA activity, respectively. Gray squares of the heatmap indicate absence of expression. (B) CMA activation score based on expression of CMA network genes in different tissues of RT under hypoxia (n=6). Data represent mean ± SEM (Two-way ANOVA). T= tissue and H= hypoxia.

## DISCUSSION

While extensively documented in the current literature, CMA is primarily investigated as a potential therapeutic target^12^. However, recent studies opened up new avenues regarding the presence of this pathway in species beyond those traditionally investigated in biomedical research^16,17^. The role of CMA in maintaining cellular homeostasis supports the idea that it could also be decisive for species exposed to numerous threats and a changing environment. Throughout this study, we have presented compelling evidence that supports the existence and crucial role of CMA in RT.

Firstly, we found that most genes of the CMA network are gradually activated throughout the development of the RT embryo. Then, quantification of the transcript levels of both CMA rate-limiting factors *lamp2a* paralogs revealed their presence in various tissues, especially those involved in metabolism, and a high heterogeneity of their expression levels, with *lamp2a* from C14 being very weakly expressed compared to its counterpart on C31. Interestingly, the low expression level of the former paralog associated with its apparent non or limited involvement in CMA activity^17^, suggests that it is under ongoing pseudogenization, as more than half of the genes duplicated during the 4th round of WGD (100 million years ago) that occurred in the salmonids ancestor^42^. Another noteworthy observation is the induction of the expression of both *lamp2a* paralogs by fasting. In mammals, although nutrient deprivation is a potent activator of CMA, it relies on reduced degradation of LAMP2A and relocation of this receptor at the lysosomal membrane^43^ and not on the control of *LAMP2A* transcription. Overall, these data highlight general conservation of the CMA machinery in RT, but also show differences with mammals in the mechanisms involved in its regulation, which remain to be further investigated.

To demonstrate CMA activity in RT, we then isolated CMA-active lysosomes from liver samples and established their ability to bind and translocate a CMA substrate. To our knowledge, this is the first *in vivo* evidence for the existence of CMA activity (or CMA-like activity) in a non-mammalian organism. This finding is in line with the recent demonstration of CMA activity in medaka cell lines using the CMA reporter KFERQ-mCherry1^16^ and provides clear evidence of that the CMA mechanism is also functional in fish. Not only does the binding and uptake of GAPDH show the capability of RT lysosomes to carry out CMA, it also demonstrates their capacity to recognize a KFERQ-like motif. This finding raises new questions regarding the KFERQ-like motif conservation throughout vertebrate evolution and whether new CMA substrates are introduced in other species therefore diversifying the scope of influence of CMA.

To advance further into unraveling the physiological role of CMA in RT, we generated a unique RT line KO for *lamp2a* C31, as our latest in cell-based knockdown experiments show that Lamp2A C14 has minimal, if any, contribution to total CMA activity^17^. As expected, the mutant fish showed a total loss of C31 *lamp2a* mRNA expression. However, they also exhibited tissue-specific modulations of the expression of the other two *lamp2* splice variant transcripts (namely, C31 *lamp2b* and C31 *lamp2c*) compared to WT. Although encoded by the same gene, previous studies reported distinct regulatory patterns for the 3 isoforms in response to various situations^30, 44, 45, 46, 47^, providing evidence for the presence of post-transcriptional mechanisms that govern the expression of *lamp2* gene(s). These mechanisms may underlie the observed variations in the expression of *lamp2b* and *lamp2c* in L2AC31-KO fish and warrant further investigation. More surprising was the induction of C14 *lamp2a* mRNA expression (carried by a different gene) in mutant RTs, which support the existence of compensatory mechanisms whose molecular determinants remain to be determined.

At a phenotypic level, mutant fish presented increased growth, enlarged livers, greater visceral weights, and a trend of lower postprandial blood glucose levels, hinting at metabolic disturbances. By investigating the state of the proteome upon *lamp2a* inactivation, we obtained a closer glimpse at CMA’s involvement in the regulation of the RT’s metabolism. Overall, Gene Ontology and network analysis highlighted clusters related to metabolic processes, mitochondrial activity, antioxidant functions and proteolytic pathways, providing compelling evidence that CMA plays a critical role as a gatekeeper of hepatic proteostasis in RT. Interestingly, several metabolic enzymes identified as *bona fide* CMA substrates (e.g., ALDOA and MDH) in mice^35^ were up regulated upon lamp2a inactivation in RT, suggesting a conservation of at least some CMA targets across vertebrates. In this sense, it is worth noting that these enzymes present KFERQ-like motifs in RT as well. Looking ahead, it will be of major interest to determine precisely which proteins are specifically targeted by CMA in this species. Another important issue will be to compare the physiological consequences of the deletion of *lamp2a* C31 with that of *lamp2a* C14 or that of the two *lamp2a*s.

As we witnessed the essential role of CMA to maintain cellular homeostasis upon an external stress, new questions arose as to the involvement of CMA in fish resilience and whether it could be harnessed as a stress gauge. To this end, we adapted and validated in RT the CMA score, as recently defined by Bourdenx et al^15^ to generate an index accounting for the CMA status. As CMA is activated in harmful situations such as oxidative stress, genotoxic damage or hypoxia^12^, we subjected RTs to moderate hypoxic conditions. In response to reduced DO levels, our investigation of multiple tissues consistently revealed a significant increase in the CMA score. This level of precision shows that CMA machinery activation is finely tuned and depends on tissue- or even cell-type, as demonstrated in mammalian cells^15^. Furthermore, it opens new perspectives regarding the potential of quickly measuring CMA status in non-invasive tissues of fish (e.g. blood or mucus) but also in uncommon species, also subjected to increasing severity of climate events, especially when unable to measure CMA activity conventionally (e.g., lysosome isolation). Nonetheless, further investigation is required (e.g., identification of specific CMA substrates, exposure of L2AC31-KO fish to different environmental stressors) in order to truly identify the mechanisms though which CMA maintains proteostasis.

## ACKNOWLEDGMENTS

We thank Cécile Heraud (INRAe) for the GSSG:GSH measurements, Christel Poujol for technical assistance at the Bordeaux Imaging Center (BIC), CNRS-INSERM and Bordeaux University, member of the national infrastructure France BioImaging and supported by the French National Research Agency (ANR-10-INBS-04). Lipidomic analyses were performed on the Bordeaux Metabolome Facility-MetaboHUB (ANR-11-INBS-0010). We thank Alexandre Stella from the ProteoToul Services - Proteomics Facility of Toulouse for the proteomics analysis. We also acknowledge technical and animal caretaking contributions from NCCCWA personal Josh Kretzer, Vanessa Panaway, and Joe Beach. Mention of trade names is solely to provide accurate information and does not reflect endorsement by the USDA. The USDA is an equal opportunity employer and provider. We would also like to thanks Lucie Marandel for kindly providing us with RT samples from her experiment^48^ and L. Peron for manufacturing the light emitting device. S.S. received a doctoral fellowship from the E2S-UPPA.

## AUTHOR CONTRIBUTIONS

I.S. and B.C. got the funding; S.S., B.C. and I.S. conceived and planned the study; S.S., K.D., V.V., I-G.P., L.M.R., E.C., B.C., F.D., P.V.D. and A.B. performed the experiments; S.S. and I.S. analyzed the data; S.S., E.J.V., K.D., V.V., I-G.P., M.G., F.B., A.H., S.F-D., B.C. and I.S. discussed the data and future experiments; S.S. and I.S. wrote the original draft of the manuscript; S.S., E.J.V., F.B., A.H., M.G., B.C. and I.S. reviewed and edited the final version of the manuscript; I.S. and B.C. supervised the project and had primary responsibility for final content.

## DECLARATION OF INTERESTS

The authors declare no competing interests.

## MATERIALS AND METHODS

Further information and requests for resources and reagents should be directed to the lead contact, Dr. Iban Seiliez

### Ethical statements

All *in vivo* experiments were conducted according to EU legal frameworks related to the protection of animals used for scientific purposes (directive 2010/63/EU), the guidelines of the French legislation governing the ethical treatment of animals (decree no. 2001-464, 29 May 2001) and/or the Institutional Animal Care and Use Committee at the United States Department of Agriculture National Center for Cool and Cold Water Aquaculture.

### Production, husbandry and treatments of L2AC31-KO fish

A commercial diet was provided for general rearing (Finfish G, Zeigler Bros, Inc., Gardners, PA), unless otherwise noted. Water flow was adjusted to maintain appropriate dissolved oxygen (> 8 ppm); water temperatures ranged from 12.5 – 13.5 °C.

Gene editing reagents were purchased from Integrated DNA Technologies (IDT) from the Alt-R CRISPR-Cas9 System product line. Online programs (crispr.wustl.edu/crispr/index.html, crispr.med.harvard.edu/sgRNAScorer/) were used to generate chromosome-specific crRNA sequences (5’-CCGTGCCCACTTTCTTTACT-3’ and 5’-AGAAGCGGCTGTGTTAACGA-3’) that cut on either side of the 9a exon of the lamp2a gene on C31, with the goal of excising the 9a exon to disrupt Lamp2a C31 protein function. Chromosome/gene specific forward primers internal to exon 9a and reverse primers within a downstream intron were used to assess whether an intact exon was present (5’-TCGGGGTTGCCTTGACTT-3’ and 5’-GGTCATCTGGATAATTGTTGATTT-3’). Two crRNAs that cut on either side of the C31 9a exons were diluted to 100 µM, as was tracrRNA, in the supplied IDTE buffer. The gRNA solution was generated by combining 1.5 µL of each crRNA, 6 µL tracrRNA, and 8.0 µL Duplex buffer. The solution was heated at 95 °C for 5 min and cooled to room temperature. Cas9 protein was diluted to 2.0 µg/µL in Cas9 working buffer (20 mM HEPES, 150 mM KCl, pH 7.5) and combined with an equal volume of gRNA solution. Ribonucleoprotein (RNP) complexes were assembled by incubating the gRNA + Cas9 mixture at 37 °C for 10 min. Phenol red was used to visualize RNP delivery.

Eggs and milt were collected from two and three-year broodstock held at the facility. Prior to fertilization, eggs were rinsed with milt activation solution (102.8 mM NaCl, 10 mM Tris, 20 mM glycine, pH 9.0, 1 mM reduced glutathione). Each fertilization lot was produced using a single female crossed with a single male. The microinjection procedure was modified from previously published procedures for RT and Atlantic salmon^49,50^. Fertilized eggs (embryos) were held in milt activation solution in an incubator at 10 °C and injected 3 – 7 h post-fertilization. Immediately prior to injection, embryos were stabilized on a tray within a Petri dish and submerged in 10 °C isotonic saline solution (155 mM NaCl, 3.22 mM KCl, 2 mM CaCl_2_). A handheld positive displacement microinjector was used to deliver between 100-200 nL of the RNP complex directly into the blastodisc, visualized using a stereomicroscope (Nikon SMZ1000). Once injected, embryos were transferred to spring water (10 – 14°C) for hatching. Injected embryos (potential mutants) and untreated embryos (controls) from the same fertilization lot were combined upon hatching for collective rearing. At approximately 20 g (∼7 month old), fish were anesthetized with tricaine methanesulfonate (MS222, 100 mg/L), adipose fin clipped for genotyping, and tagged with passive integrated transponders (PIT tags) inserted in the intraperitoneal cavity. Genotyping was completed during the grow-out period to distinguish between 1) WT controls, 2) mutants with indels at target sites but retained the 9a exon (intact gene), and 3) mutants with excised exons at the lamp2a gene on C31. Twelve fish were identified that exhibited mosaicism for excision of the 9a exon; these fish, including WT controls from the same fertilization lots were used to produce an F1 generation. A subsample of 10 sacfry from each offspring lot were harvested after hatch to screen for gene mutagenesis. Approximately 50% of the F1 offspring from a single cross exhibited complete excision of the 9a exon on the C31 gene. This cross, along with the associated F1 control cross, were hatched and reared separately until PIT tagging and fin clipping at approximately 20 g body weight. After PIT tagging, fish were comingled for grow-out under standard rearing protocols.

At approximately one year of age, a subset of 68 fish from the F1 mutant (51 fish) and F1 control (17 fish) groups were stocked into two 800 L tanks and acclimated to these conditions for two weeks while consuming a commercially available diet. Feed was withheld for 18 hours before recording weights and fork lengths on individual fish; for the next seven weeks fish were fed to satiation with a high carbohydrate diet (Table S5A). Feed was provided using automatic feeders set to dispense feed at a fixed percent of tank biomass between 08:00 h and 14:00 h, with hand feeding at 14:30 h each day. Ration was adjusted so hand feeding to satiation was approximately 5 – 10% of the total daily ration. Weights and lengths were recorded after week 4 and at the end of the study (week 7). Feed was withheld the day of sampling. At the final sampling, a subset of mutants and controls (n = 9 per group, n = 4-5 per tank) were euthanized with a lethal dose of MS222 (300 mg/L). Blood was collected from caudal vasculature into heparinized tubes and placed on ice for subsequent processing for separation of plasma and storage at -80 °C. The viscera was removed and liver and gastrointestinal tract weights were recorded. Samples of liver and distal intestine were placed in histology cassettes and submerged in 10% neutral-buffered formalin with 4% formaldehyde prior to paraffin embedding. Subsamples of liver, distal intestine, and whole brain were frozen in liquid nitrogen and stored at -80 °C.

### Other in vivo experiments

Samples used to investigate the expression of CMA-related genes during RT development were generated in a previously published study^48^. Briefly, spawns were fertilized synchronously with neomale sperm and reared in separate tanks at 8°C at the INRAE experimental facilities, Lees-Athas, France. Fish were sampled according to Vernier et al.1969^23^ at stages 2, 5, 6, 7, 8, 10, 12, 15, 22 and 23. Embryos were directly snap-frozen while juveniles anaesthetized and killed in benzocaine (60 mg/L). Fish were consistently sampled at 12:00 h for each ontogenetic stage, to avoid potential circadian effects.

Samples used to investigate the tissue-specific expression of both lamp2a genes were generated in a previously published study^51^. In details, all-female RT from a spring-spawning strain were used and kept under a natural photoperiod at 17°C at the INRAE experimental facilities in Donzacq, France. Twice a day, fish were manually fed ad libitum and considered to be satiated when only a few pellets remained on the bottom of the tank. At the end of experiment, fish were anesthetized and euthanized 6 h after the last meal. Then tissues were sampled and immediately stored at −80 °C.

For the hypoxia experiment, monosex female RTs from a spring-spawning strain were used and kept under a natural photoperiod^51^. The 12 fish studied were part of a pool of 240 fish (i.e., initial mean weight of 288 ± 9 g) randomly distributed in 12 tanks of 150 L each, for an initial mean biomass of 18 ± 1 kg/m3. A factorial experimental design was used to obtain two DO levels (normoxia vs. moderate hypoxia). The treatment was applied for four weeks.

Hypoxic conditions were created by reducing the water flow. DO concentrations were measured at the tank’s outlet twice daily using an oximeter (HatchLange HQ 40D). The mean DO concentrations in the hypoxia and normoxia tanks were 5.6 ± 0.5 and 7.7 ± 0.3 ppm, respectively. Mean DO concentrations were higher than those needed to support positive growth, feeding and swimming activities. Total ammonia nitrogen (NH4-+N), nitrite-nitrogen (NO2−-N) and nitrate-nitrogen (NO3−-N) were analyzed twice a week using a commercial kit based on colorimetric method (© Tetra). No trace of these elements was recorded. The temperature remained constant (17.0 ± 0.5 °C). We can therefore consider that the decrease in water level only caused a decrease in oxygen.

Twice a day, fish were manually fed ad libitum and considered to be satiated when only a few pellets remained on the bottom of the tank.

### RTgutGC cell line

For in vitro experiments, the RTgutGC cell line^52^ derived from RT was routinely grown in Leibovitz’s L-15 medium supplemented with 10% fetal bovine serum (FBS, #10270-106), 2 mM sodium pyruvate (NaPyr, #11360-070), 100 units/mL penicillin and 100 g/mL streptomycin (#14065-056), all provided by Gibco (Thermo Fisher scientific) and 25 mM HEPES (#BP299-1, Fisher Bioreagents, Fisher Scientific SAS, Illkirch Graffenstaden, France). Cells were maintained at 18 °C, the medium was replaced twice a week and cells were passaged at 80% of confluence. Cell counting was achieved using a Cellometer K2 (Nexcelom Bioscience LLC, Lawrence, MA, USA) to plate cells prior to the experiments. Cells were seeded at a density of 50-60% for RNA isolation. Prior to the treatments, cells were washed twice with PBS before the exposure to the appropriate treatment. The treatments consisted in applying a mild-oxidative stress using CT medium supplemented with hydrogen peroxide (H2O2 25 µM) or using a Hank’s balanced salt solution (HBSS) (#14025-092, Gibco) supplemented with 25 mM HEPES as a starvation medium.

Nucleofector 2b Device (Lonza, Colmar, France) and the Cell line Nucleofector Kit T (Lonza, VCA-1002) were used for the plasmids cells transfections (1-5 µg). RTgutGC cells stably-transfected with the KFERQ-PA-mCherry1 construction were selected for the resistance to the selective geneticin antibiotic (G418; Gibco, 11811) before cell sorting using a FACS Aria 2-Blue 6-Violet 3-Red 5-YelGr, 2 UV laser configuration (BD Biosciences, Le Pont de Claix Cedex, France) in biosafety cabinet.

### RNA extraction and quantitative RT-PCR analysis

To extract total RNA from individual samples, tissues were ground in TRIzol reagent (Invitrogen, Carlsbad, California, USA). 50-100 mg of tissue per mL of reagent were sampled and homogenized in a Precellys tissue homogenizer (Bertin Technologies, Montigny le Bretonneux, France). The Super-Script III RNAse H-Reverse transcriptase kit (Invitrogen) was used with random primers (Promega, Charbonniéres, France) to synthesize cDNA from 1μg of total RNA. The primer sequences used are listed in Tables S1 and S6. Primers were validated by testing their efficiency on pooled cDNA and subsequently sequencing the amplified products. A Roche Lightcycler 480 system (Roche Diagnostics, Neuilly-sur-Seine, France) was used for real-time RT-PCR following the protocol from Plagnes et al. 2008^53^. Quadruplicates were analyzed and standardized with luciferase (Promega) or stably expressed housekeeping genes (18S, or *β*-actin). The E-method^54^ was used for the developmental and tissue-specific expression analysis, and the ΔCT method^55^ for the expression of target genes in KO-RTs and CMA score calculation.

### *In situ* hybridization

The liver and distal portion of the intestine of starved fish (48h) and fish fed with a commercial diet were sampled and immediately fixed with 4% paraformaldehyde (PFA 4%). Samples were then rinsed in PBS, dehydrated through gradual ethanol and butanol baths, embedded in paraffin and cut in 7μM transversal sections. The sections were placed on superfrost adhesion slides (VWR®). The following steps were performed according to the BaseScope™ Detection Reagent Kit v2 – RED User Manual. Samples were deparaffinized and dried after a series of xylene and alcohol baths. To facilitate target access and detection sensitivity, we applied a target retrieval, then a hydrophobic barrier was set and a tissue-specific protease applied. Both BASEscope lamp2a probes, designed by ACD bio® were applied before the amplification phase during which multiple probes are hybridized to amplify the signal. A fast red substrate was then added to visualize the target RNA. A nanozoomer - slides scanner (Hamamatsu NANOZOOMER 2.0HT) was used to visualize whole tissue sections. To quantify the mRNA signal in the liver, equal size images of hepatocyte-covered area were used to count the particles (mRNA). However, as the signal was not evenly distributed in the distal intestine, each image was calibrated using ImageJ^56^ so as to only quantify the signal in epithelium area.

### Lysosome isolation and GAPDH binding/uptake assay

The lysosome isolation protocol^26^ was adapted to RT, as it is originally targeted to rat liver. A piece of starved fish liver was sampled (∼1-1.5 g), washed with cold PBS, minced with a clean scalpel and homogenized in a conical tube. Some of the homogenate was put aside as a marker for total sample proteins (TLH) and the rest was filtered through a cotton gauze. After a series of centrifugation, a mix of mitochondria and the lysosomal fraction was collected. A discontinuous Nycodenz® (Proteogenix) gradient was then generated and ultracentrifuged (141000 g for one hour at 4°C) to create different interphases with specific content. From bottom to top were isolated mitochondria, a mixture of mitochondria and lysosomes, a mix population of lysosomes and lysosomes with high CMA activity. Those phases were collected with a Pasteur pipet and washed by centrifugation. The third interphase was resuspended and centrifuged to only be able to collect high CMA activity lysosomes.

For the GAPDH binding/uptake assay, protein concentrations were measured by Qubit assay (Thermo Fisher Scientific). Isolated lysosomes were incubated in the uptake assay buffer: 300 mM sucrose, 10 mM MOPS, pH 7.2, 10 mM ATP (Sigma A2383), 0.01 μg/μL biologically active HSC70 (AbCam, ab78431). For the conditions needing a CMA substrate, samples were incubated with 0.025 μg/μL GAPDH (LP00008). In conditions involving PK, samples were incubated with 1 μL of PK (Sigma P6556). To prevent lysosomal degradation, chymostatin (Sigma C7268) was used. All samples were centrifuged and the pellet was then washed and prepared for SDS-PAGE and western blotting.

To measure lysosomal integrity, isolated lysosomes were diluted in a buffer of 300mM sucrose, 10 mM MOPS, pH 7.2, and 10 mM ATP. Then the sample was centrifuged to separate the pellet (intact lysosomes) from the supernatant (broken lysosomes). Lysosomal integrity was measured by the β-hexosaminidase assay with the fluorogenic substrate 4-methylumbelliferyl-2-acetamido-2-deoxy-β-D-glucopyranoside (MUG) and and only experiments with >90% intact lysosomes were considered.

### Western blotting analysis

To verify if RT lysosomes display known characteristics of CMA+ lysosomes, the isolated fractions (CMA+ lysosomes, CMA+/- lysosomes, lysosomes/mitochondria and mitochondria) were subjected to SDS-PAGE and immunoblotted using the following antibodies: LAMP1 (ab24170), HSC70 (AS05 062) and TIM23 (BD611222). The total tissue homogenate was also quantified as it contains the biological components of all fractions (TLH). To quantify GAPDH, GAPDH (ab8245) was used. We used LC3B (CST3868S) to measure steady-state levels of autophagosome component LC3. After being washed, membranes were incubated with corresponding secondary antibodies such as: HRP anti-rabbit (926-80011) HRP anti-mouse (926-80010). Bands could be visualized by using the Invitrogen™ iBright™ FL1500 Imaging System and analyzed with the iBright analysis software (version 5.0). To ensure protein normalization, a No-Stain Protein Labeling Reagent (A44717) and β Actin (sc-47778) were used.

### Plasma glucose levels

Blood glucose measurements were analyzed immediately upon collection using a Prodigy AutoCode glucometer (Prodigy Diabetes Care, LLC, Charlotte, NC, USA).

### Glycogen content

Glycogen content of WT and L2AC31-KO RT was analyzed on lyophilized liver through a hydrolysis technique described by Good et al. (1933). Each sample was homogenized in 1 mol.L-1 HCl (VWR, United States) and free glucose content was measured. Another measurement of free glucose occurred after a 10min centrifugation at 10.000 g using the Amplite Fluorimetric Glucose Quantification Kit (AAT Bioquest, Inc., United States) according to the manufacturer’s instructions. The remainder ground tissue was boiled for 2.5 h at 100 °C before being neutralized with 5 mol.L-1 KOH (VWR, United States). Total glucose (free glucose + glucose produced by glycogen hydrolysis) was measured then free glucose levels were subtracted to determine the glycogen content.

### Lipid content

Lipids were extracted and purified according to Folch et al.1957^57^ and Alannan et al.2022^58^. Briefly, the lipids from about 100 mg of fresh weight were homogenized in an Eppendorf tube with 2 x 1 mL of ice-cold isopropanol with a TissueLyser. The homogenates were transferred in a glass tube and 1 mL of ice-cold isopropanol was used to rinse the Eppendorf tube. The samples were then placed at 85 °C for 30 min to inactive lipases. After letting the samples cool down, lipids were extracted with 3 mL of choloroform/methanol 2:1 for 2 h at room temperature, with 3 mL of chloroform overnight at 4 °C and then with 2 mL for an additional 4 h at room temperature. Between each extraction, the samples were centrifuged for 5 min at 3,500 g and the organic phase was collected and transferred to a fresh tube where all organic phases were ultimately pooled (final volume = 11 mL). Polar contaminants such as proteins or nucleic acids were removed by adding 4 mL of NaCl 0.9 % 100 mM Tris and shaking vigorously. After phase separation by centrifugation (15 min at 3.500 g) the organic phase was collected and the solvent was evaporated. The lipids were then resuspended in chloroform:methanol (1:1, v/v) for a final concentration of 0,1 mg equivalent fresh weight/µl. For quantification of neutral lipids, 50 µL (= 5 mg fresh weight) of total lipids were applied onto a silica-coated chromatography plate (Silicagel 60 F254, 10x 20 cm; Merck, Rahway, NJ) and developed with hexane/ethyl ether/acetic acid (21. 25:3.75:0.25, v/v) The plates were then immersed in a copper acetate solution (3% copper acid + 8 % phosphoric acid in distilled water) and heated at 115 °C for 30 min. Lipids were identified by co-migration with known standards and quantified by densitometry analysis using a TLC scanner 3 (CAMAG, Muttenz, Switzerland). Fatty acid methyl esters were obtained by transmethylation at 85 °C for 1 h with 0.5 M sulfuric acid in methanol containing 2 % (v/v) dimethoxypropane and 5 µg/mL of heptadecanoic acid (C17:0) as internal standards. After cooling, 1 mL of NaCl (2.5 %, w/v) was added, and fatty acyl chains were extracted with 1 mL hexane. GC-FID was performed using an Agilent 7890 gas chromatograph equipped with a DB-WAX column (15 m x 0.53 mm, 1 µm; Agilent) and flame ionization detection. The temperature gradient was 160 °C for 1 min, increased to 190 °C at 20 °C/min, increased to 210 °C at 5 °C/min and then remained at 210 °C for 5 min. FAMES were identified by comparing their retention times with commercial fatty acid standards (Sigma-Aldrich) and quantified using ChemStation (Agilent) to calculate the peak surfaces, and then comparing them with the C17:0 response

### Quantification of GSH, GSSG in liver

AccQ·Fluor™ borate buffer was purchased from Waters® (Massachusetts, USA). Glutathione reduced (GSH), Glutathione oxidized (GSSG), EthyleneDiamineTetraacetic Acid (EDTA), Phosphoric acid (H3PO4), Potassium dihydrogen phosphate (KH2PO4) and MetaPhosphoric Acid (MPA) were purchased from Sigma-Aldrich® (Germany). Other reagents that used were of HPLC grade. All organic solvents used were gradient HPLC grade (ADL & Prochilab, Lormont, France). Ultrapure water was daily made with a MilliQ Direct8 system (Millipore, Bedford, MA, USA).Each sample (almost exactly 50 mg) was homogenized with Precellys® tissue homogenizer in 300 µL of 20 mM phosphate buffer, 1 mM EDTA (pH = 6.5 ± 0.05). After centrifugation (14.000 g, 20 min, 4 °C), deproteinization of the supernatant was performed with a volume-to-volume 10 % metaphosphoric acid solution. After centrifugation (14.000 g, 5 min, 4 °C), the supernatant was filtered with a 0.22 µm PVDF unit. Metabolite mixtures were stored at -20°C (no more than 1 week) until LC-UV analysis.Chromatographic separation was achieved on Waters® Symmetry Shield RP18 column (4.6 mm × 150 mm i.d. 3.5 μm). The column was operated at 30 °C. The injection volume was 50 μL and the flow rate was set at 0.7 mL/min. A ternary solvent system was used, consisting of (A) pH2.7 20 mM phosphate buffer, (B) acetonitrile, (C) methanol. The mobile phase was filtered through in-line 0.2 μm membrane filters. The following gradient elution was employed: 0–10 min : 99.5 % A, 0 % B; 10.5 min : 40 % A, 60 % B; 10-13 min : 40 % A, 60 % B; 13.5 min : 99.5 % A, 0.5 % B ; 13.5 – 20 min (column equilibration) : 99.5 % A, 0.5 % B. The eluate was monitored with absorbance detection at 210 nm during the whole run. Waters® Empower™ Pro software was used for data acquisition. Metabolites were identified comparing their Retention Time (RT) to standard ones. Quantification was based on integration of peak areas and compared to the standard calibration curves (R2 correlation > 0.999) of each metabolite of interest (6 biological levels, 3 repetitions). Calibration curves were linear from 0.2 to ∼1 000 pmol/injection for GSSG and from 2 to ∼10 000 pmol/injection for GSH.

### Sample preparation for label-free proteomics analysis

Proteins from WT and KO25-trouts were extracted using the Thermo Scientific™ T-PER™ Tissue Protein Extraction Reagent (ref 78510). Protein concentrations were then measured by bicinchoninic acid (BCA) assay (Interchim; ref UP95424A). For each sample 50 µg of dried protein extracts were solubilized with 25 µl of 5% SDS. Proteins were submitted to reduction and alkylation of cysteine residues by addition of TCEP and chloroacetamide to a final concentration respectively of 10 mM and 40 mM. Protein samples were then processed for trypsin digestion on S-trap Micro devices (Protifi) according to manufacturer’s protocol, with the following modifications: precipitation was performed using 216 µl S-Trap buffer, 4 µg trypsin was added per sample for digestion in 25 µL ammonium bicarbonate 50 mM.

### NanoLC-MS/MS analysis of proteins

Tryptic peptides were resuspended in 35 µL of 2 % acetonitrile and 0.05 % trifluoroacetic acid and analyzed by nano-liquid chromatography (LC) coupled to tandem MS, using an UltiMate 3000 system (NCS-3500RS Nano/Cap System; Thermo Fisher Scientific) coupled to an Orbitrap Exploris 480 mass spectrometer equipped with a FAIMS Pro device (Thermo Fisher Scientific). 1 µg of each sample was injected into the analytical C18 column (75 µm inner diameter × 50 cm, Acclaim PepMap 2 µm C18 Thermo Fisher Scientific) equilibrated in 97.5 % solvent A (5 % acetonitrile, 0.2 % formic acid) and 2.5 % solvent B (80 % acetonitrile, 0.2 % formic acid). Peptides were eluted using a 2.5 % - 40 % gradient of solvent B over 62 min at a flow rate of 300 nL/min. The mass spectrometer was operated in data-dependent acquisition mode with the Xcalibur software. MS survey scans were acquired with a resolution of 60.000 and a normalized AGC target of 300 %. Two compensation voltages were applied (-45 v /-60 v). For 0.8 s most intense ions were selected for fragmentation by high-energy collision-induced dissociation, and the resulting fragments were analyzed at a resolution of 30.000, using a normalized AGC target of 100 %. Dynamic exclusion was used within 45s to prevent repetitive selection of the same peptide.

### Bioinformatics analysis of mass spectrometry raw files

Raw MS files were processed with the Mascot software (version 2.7.0) for database search and Proline55 for label-free quantitative analysis (version 2.1.2). Data were searched against Rainbow trout entries of Uniprot protein database (46.650 sequences; 17.861.404 residues). Carbamidomethylation of cysteines was set as a fixed modification, whereas oxidation of methionine was set as variable modifications. Specificity of trypsin/P digestion was set for cleavage after K or R, and two missed trypsin cleavage sites were allowed. The mass tolerance was set to 10 ppm for the precursor and to 20 mmu in tandem MS mode. Minimum peptide length was set to 7 amino acids, and identification results were further validated in Proline by the target decoy approach using a reverse database at both a PSM and protein false-discovery rate of 1%. For label-free relative quantification of the proteins across biological replicates and conditions, cross-assignment of peptide ions peaks was enabled inside group with a match time window of 1 min, after alignment of the runs with a tolerance of +/- 600 s. Median Ratio Fitting computes a matrix of abundance ratios calculated between any two runs from ion abundances for each protein. For each pair-wise ratio, the median of the ion ratios is then calculated and used to represent the protein ratio between these two runs. A least-squares regression is then performed to approximate the relative abundance of the protein in each run in the dataset. This abundance is finally rescaled to the sum of the ion abundances across runs. A Student t-test (two-tailed t-test, equal variances) was then performed on log2 transformed values to analyze differences in protein abundance in all biologic group comparisons. Significance level was set at p = 0.05, and ratios were considered relevant if higher than +/- 2. The results were ranked based on fold change (>1.41) and p-value (<0.05) of t-test comparing WT and L2AC31-KO groups. Gene Ontology enrichment maps were generated in CytoScape (3.9.1) using EnrichmentMAP (3.3.5) plugin, and then annotated with AutoAnnotate (1.4.0) app. The presence of KFERQ motifs within the protein sequences was assessed using the KFERQ finder app V0.8^13^. Furthermore, an ontology analysis was conducted using Enrichr^59^. Clusters are grouped by major functional association.

### CMA score

Each gene of the CMA network is attributed a weight. As some genes are present in multiple copies (paralogs) in RT, each paralog was attributed a weight of 1/n, with n copies. As LAMP2A is the rate-limiting factor in CMA, it was given a weight of 2 (1/paralog). Weight values for LAMP2A copies were measured with the relative contribution of each paralog to CMA activity in a RT cell line stably expressing the KFERQ-PA-mCherry1 reporter^17^. Furthermore, a modulating score (+1 or -1) was applied to the values, whether the gene has a positive or negative impact on CMA. Finally, a sum of all values was calculated to obtain a final score.

### Quantification and Statistical analysis

Data is reported as means + SEM, percentages + SEM. Normality assumptions were tested before conducting statistical tests and parametric tests conducted only when they were met. Differences between more than two groups were evaluated by one-way ANOVA followed by Tukey’s multiple comparison (same sample size) post-hoc test. In the case of non-parametric tests, Kruskal-Wallis or Friedman test followed by Dunn’s multiple comparisons were conducted. When comparing two groups we used a parametric two-tailed unpaired Student’s T-test, or a non-parametric Mann Whitney test. All statistical analyses were performed using GraphPad Prism version 8.0.1 for Windows (GraphPad Software, Inc., www.graphpad.com) and a *p*-value < 0.05 was set as a level of significance. All data generated in this study and presented in the figures are provided along with their statistic report in the Dataset S1.

## SUPPLEMENTAL INFORMATION

**Table S1.** CMA score network genes and corresponding primers (forward and reverse).

**Table S2.** Up-regulated proteins in L2AC31-KO vs WT RTs. A Student t-test (two-tailed t-test, equal variances) was performed on log2 transformed values to analyze differences in protein abundance. The results were ranked based on fold change (>1.41) and p-value (<0.05) of t-test comparing WT and L2AC31-KO groups.

**Table S3.** Down-regulated proteins in L2AC31-KO vs WT RTs. A Student t-test (two-tailed t-test, equal variances) was performed on log2 transformed values to analyze differences in protein abundance. The results were ranked based on fold change (>1.41) and p-value (<0.05) of t-test comparing WT and L2AC31-KO groups.

**Table S4.** Occurrence of KFERQ-like motifs in differentially-regulated proteins in L2AC31-KO RTs. KFERQ-like motifs were searched using the KFERQ finder app V0.8^13^ and are categorized as canonical (canon.), phosphorylated (phos.) or acetylated (acet.).

**Table S5.** Formulation and proximate composition of the experimental diet used in the knock-out experiment (High CHO diet). HighCHO: high-carbohydrate diet. ***1*** Sopropeche, Boulogne-sur-Mer, France. ***2*** Gelatinized corn starch; Roquette, Lestrem, France. ***3*** Sopropeche, Boulogne-sur-Mer, France.***4*** Louis François, Marne-la-Vallée, France. ***5*** Supplied the following (/kg diet): DL-a tocopherol acetate 60 IU, sodium menadione bisulphate 5 mg, retinyl acetate 15,000 IU, DLcholecalciferol 3,000 IU, thiamin 15 mg, riboflavin 30 mg, pyridoxine 15 mg, vit. B12 0.05 mg, nicotinic acid 175 mg, folic acid 500 mg, inositol 1,000 mg, biotin 2.5 mg, calcium panthotenate 50 mg, choline chloride 2000 mg. ***6*** Supplied the following (/kg diet): calcium carbonate (40% Ca) 2.15 g, magnesium oxide (60% Mg) 1.24 g, ferric citrate 0.2 g, potassium iodide (75% I) 0.4 mg, zinc sulphate (36% Zn) 0.4 g, copper sulphate (25% Cu) 0.3 g, manganese sulphate (33% Mn) 0.3 g, dibasic calcium phosphate (20% Ca, 18% P) 5 g, cobalt sulphate 2 mg, sodium selenite (30% Se) 3 mg, potassium chloride 0.9 g, sodium chloride 0.4 g.

**Table S6.** List of other primers used in study (forward and reverse).

**Figure S1. mRNA expression of *lamp2b* and *lamp2c* splice variants in WT *vs* L2AC31-KO RTs. (A)** Sequence alignment of *lamp2a* in WT vs L2AC31-KO RT showing the loss of exon A (highlighted in yellow) and the CRISPR targets (red font). (B) and (C) show the mRNA expression of *lamp2b* from C31 and C14, respectively. (D) and (E) show the mRNA expression of *lamp2c* from C31 and C14, respectively. Values are mean + SEM (n=7) shown by fold induction respect to WT group. T-test was used to compare between groups (*, *p*<0.05; **, *p*<0.01).

**Figure S2. *In vitro* validation of the CMA score using KFERQ-PA-mCherry1 reporter in RTgutGC cells.** (A) Graphical representation of the CMA network. Each group of proteins is based on function (effectors and modulators) and localization (lysosomal and extra-lysosomal) (modified from^15^). (B) Representative images of RT enterocytes cells visualized by fluorescence microscopy (scale bars, 20 μm) and zoomed image with scale bars of 5 μm. Cells were transfected with a photoactivable (PA) KFERQ-PA-mCherry1 fluorescent reporter (red) and then photoactivated for 10 min. Cells were untreated (Control), exposed to a mild-oxidative stress (H2O2, 25 µM) or maintained in the absence of serum (starvation) for 16 h. CMA activity was measured as the total number of KFERQ-PA-mCherry1 fluorescent puncta normalized for the number of cells after 4, 8, 16, 24 or 48 h exposure of RT enterocytes to mild-oxidative stress (C) or starvation (D) and compared with control cells using Mann-Whitney tests (*, *p*<0.05; **, *p*<0.01; ***, *p*<0.001; #, *p*<0.0001) (n=3 experiments, values are means +/- SEM). (E) Heatmap representing expression of CMA network genes in mild-oxidative stress and starvation conditions. mRNA levels were standardized with housekeeping gene 18S. (n=3, values are mean). (F) CMA score based on expression of CMA network genes under both conditions. (n=3, all values shown min. to max.). A t-test was performed comparing the control condition with each treatment condition (*, *p*<0.05).

## Notes

### Competing Interest Statement

The authors have declared no competing interest.

