## Supplementary material for "Chaperone-Mediated Autophagy in Fish: A Key Function Amid a Changing Environment": SUP FIGURES

FigS1

**A**

```
       WT   ...TTAGCTGCAATAAATGGGAATGGAAATTGTGCATAAACTGGCGTGGACCATTTGTAATATTTTACAATATTATTGGTCCCCCGGACATTTTAGTAAA
L2AC31-KO   ...TTAGCTGCAATAAATGGGAATGGAAATTGTGCATAAACTGGCGTGGACCATTTGTAATATTTTACAATATTATTGGTCCCCCGGACATTTTAGTAAA
               *****************************************************************************************************

       WT   TCCCTCCTTTGTTTTCCAACTATTCTTGACTGAGAAGTGTCTATGATTGACCATGCATTCACCATTTTGATGTTTTACCTTTCTTCCACCCTGCAACCCC
L2AC31-KO   TCCCTCCTTTGTTTTCCAACTATTCTTGACTGAGAAGTGTCTATGATTGACCATGCATTCACCATTTTGATGTTTTACCTTTCTTCCACCCTGCAACCCC
            ****************************************************************************************************

       WT   CCCGTGCCCACTTTCTTTACTGTCATCCAGCGGAGGATTGCCAAGACGATACGACAGAGAGCTGGCTTGTTCCTATAGCGGTCGGGGTTGCCTTGACTTT
L2AC31-KO   CCCGTG----------------------------------------------------------------------------------------------
            ******

       WT   ACTGGTCCTCATTGTGTTGGTTGCCTATTTCATTGGAAGAAAGCGAAACCAGGGCACTGGCTATGAGCACTTCTAAATTATCTTCACTATGCTGAGGCTA
L2AC31-KO   ----------------------------------------------------------------------------------------------------

       WT   TAACTTCGGTCATCTGGATAATTGTTGATTTAATTTGACTAATATACTGTGCAGAGTTTACTTAATGGTGAAATTCACCGGATGGAAAGCTCTTTCTCAT
L2AC31-KO   ----------------------------------------------------------------------------------------------------

       WT   GAAACAATGACTTGGAGAAGCGGCTGTGTTAACGATGGCATGTAACAGCTGTGTGTGTAATTAGAATTTATTGATTTTGATTGACACATCTTATATTCCT
L2AC31-KO   ----------------------------TTAACGATGGCATGTAACAGCTGTGTGTGTAATTAGAATTTATTGATTTTGATTGACACATCTTATATTCCT
                                        ************************************************************************

       WT   CATCACATGGTTGACTATAGATGGCTGATGACCAAATGTTTTCCTAATTGCAATTAATGCCCATGCTCTGACGGACCCTGATGAATATGTCTTGGTT...
L2AC31-KO   CATCACATGGTTGACTATAGATGGCTGATGACCAAATGTTTTCCTAATTGCAATTAATGCCAATGCTCTGACGGACCCTGATGAATATGTCTTGGTT...
            *************************************************************************************************
```

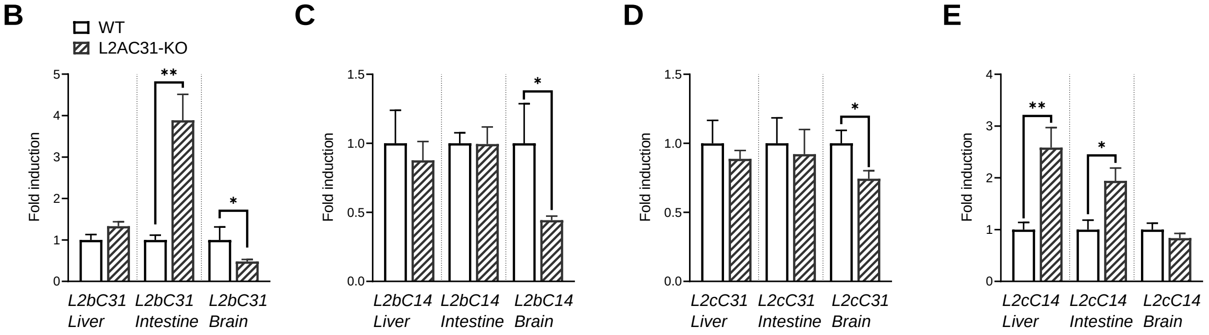

Fig S2

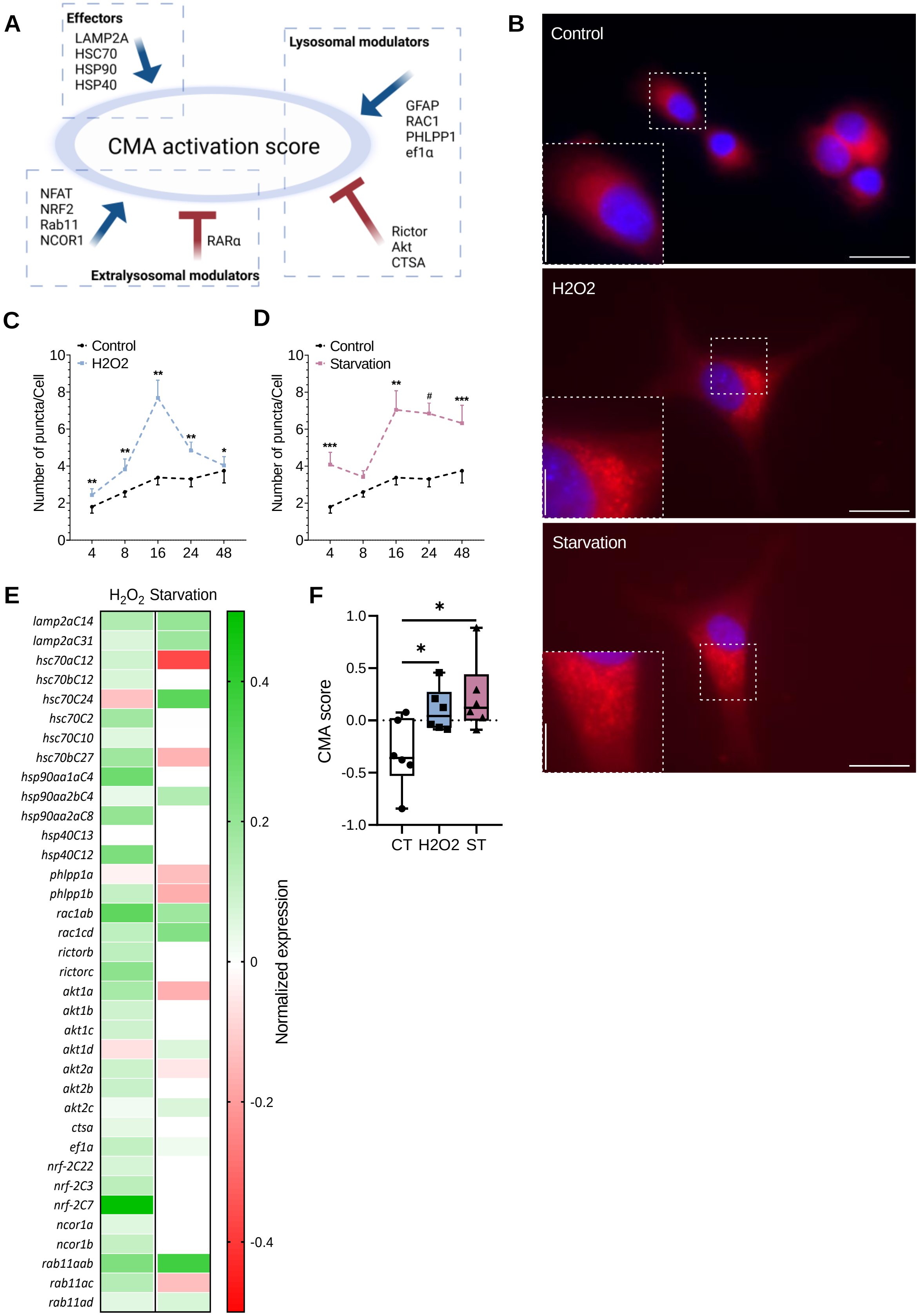
A
Effectors
LAMP2A
HSC70
HSP90
HSP40
Lysosomal modulators
GFAP
RAC1
PHLPP1
ef1α
CMA activation score
NFAT
NRF2
Rab11
NCOR1
RARα
Rictor
Akt
CTSA
Extralysosomal modulators
B
Control
H2O2
Starvation
C
Control
H2O2
Number of puncta/Cell
D
Control
Starvation
E
H2O2 Starvation
lamp2aC14
lamp2aC31
hsc70aC12
hsc70bC12
hsc70C24
hsc70C2
hsc70C10
hsc70bC27
hsp90aa1aC4
hsp90aa2bC4
hsp90aa2aC8
hsp40C13
hsp40C12
phlpp1a
phlpp1b
rac1ab
rac1cd
rictorb
rictorc
akt1a
akt1b
akt1c
akt1d
akt2a
akt2b
akt2c
ctsa
ef1a
nrf-2C22
nrf-2C3
nrf-2C7
ncor1a
ncor1b
rab11aab
rab11ac
rab11ad
Normalized expression
F
CMA score
CT H2O2 ST
