## Supplementary material for "Chaperone-Mediated Autophagy in Fish: A Key Function Amid a Changing Environment": SUP TABLES

**Table S1.** CMA network genes and corresponding primers

| Role | Name of gene | Forward sequence | Reverse sequence |
|---|---|---|---|
| Effectors | *lamp2a C14* | CAACAGGGAGCAGACATTAACT | CAATGAGGATCAGCACAGCC |
| | *lamp2a C31* | CAACAAGGAGCAGACCTTAAAC | CCCTGGTTTCGCTTTCTTCC |
| | *hspa8 C10* | TGCGTCCATTATATCCAGCA | CAAACGCTTAGCATCGAACA |
| | *hsc70a C12* | ACCTCCCCCTAACAAGCAAA | AGGCATTGTGACAAAGGCAG |
| | *hsc70b C12* | GCCCAATCTGTAGTAAAGCCAAG | CTCCAGGACTCAAAATGTAGACAAA |
| | *hspa8b C27* | ATGGCTTGGAATGGACTGAA | CATTTATTGGCTAAACTGAAAACG |
| | *hsc70 C24* | TGCTGAAAGCAACCAATCAA | TGCAGTACAAAGGCAAAAACA |
| | *hsc70 C2* | GGCTCTTCCTCTGGTCCAAC | GCTGTAGGAGTCTGCCGTAG |
| | *hsp90aa1a C4* | GAGGATTAGGACCAGTGAAATACA | TGGCGTTTTCTGATCACTGT |
| | *hsp90aa2b C4* | GAGAAGAAAGATGGGGAAGGAGAG | CTTGTCCCCACATGCGCCGTCA |
| | *hsp90aa2a C8* | GAGGGAGACGAGGACACATC | GAACAAACATCAAAGTCATCCTG |
| | *hsp90aa1b C8* | TTTTCATGACTGCTGAGTCG | TTGAATAGGGTTGAAAACAGCA |
| | *dnajb1a C13* | TCAAGCGTTACTTGACAGATACTTG | TCAGCTCCAGTGGACTTGTTT |
| | *dnajb1a C12* | TCAAGCGTTACTTGGCAGAC | TCAGCTCCAGTGGACTTGTTT |
| Modulators | *gfapab* | GCTCCGTGGAACAAACGAGT | GATGACCCACCGTATCCTGG |
| | *phlpp1a* | AACAGGTTCGGCAGAGGC | TGGGGCTTCCCTGCGTAA |
| | *phlpp1b* | TTGAAAACAGGGTCGGGGG | TGTGGGGCTTTCCTGCGTAA |
| | *rac1ab* | GTACATCCCCACAGTGTTTGA | ATCCCAAAGGCCCAAGTTGA |
| | *rac1cd* | ATGGCAAAAGAGATTGGAGCAG | CTTCCTCTTCTTGACGGGCG |
| | *rictora* | GTCCATCTCTATGCCTTGGTTT | CTGTTGGAAGTAGGGCGTGT |
| | *rictorb* | ATCATCTGTGAACTTGCTGTGA | TAGTGAAAGTCTGTGTACGGGG |
| | *rictorc* | CCCAGTAAGTCTCAGCCCAGTATG | CGTCCGACAGGTCACAGTTC |
| | *akt1a* | ACCAAGCCCAAACTCAAAGTG | GTGTGCTACTTCATCTTTTGCC |
| | *akt1b* | CGACGGTACATTCATTGGCT | AGCTGACATTCTGGTGCTACT |
| | *akt1c* | CCACAAGATGTTGACGCGTTCG | CTTTCGTCCATTCTTCCCTCTCC |
| | *akt1d* | CATTGAGGCCGTAGCAGACAG | TTCAAAGTCGTGCATAGTCACTT |

| | | | |
|---|---|---|---|
| | *akt2a* | AAAGGACCCCAAGCAGAGGTTA | AAGTAGCGGGTGTCGGTC |
| | *akt2b* | GATCCCAAGCAAAGGTTAGGG | TGCCACAGTCCAGGTTGGTAT |
| | *akt2c* | TGCCAAAGACGTGATGACACAG | TGAGTCTAAGCTGTCATACTTGTCC |
| | *akt2d* | GATGCCCCACGAGGAGAGTA | GTAGGCAGCGGATGACAAAC |
| | *ctsa* | GTTTCTCCCTGGGCTTCAAA | GACATGCCATCGTTCTGAATCAAG |
| | *ef1a* | TCCTCTTGGTCGTTTCGCTG | ACCCGAGGGACATCCTGTG |
| | *nfatc1a* | TACCTTTCCCCAAACGTCCC | GATAAACCCCGACCAGGCAG |
| | *nfatc1b* | AGGACTCCTTCAAGTCTACCGC | AATGAACAGACAGCCCCTGAG |
| | *nfatc1c* | AAGGACTCCTGCAAGTCTACAT | CGCTGGGGACACACCTGA |
| | *nfatc1d* | CAAGGACTCCTCCAAATCTACATC | CCAGAACAATGCTTGGGACA |
| | *nrf2 C22* | CTAGCACCCAACAGCAGTCA | TCAGACATTGCTGCAGCTCA |
| | *nrf2 C3* | CCTGTGGACGACTTCAACGA | AGCCACCTTATTCTTGCCCC |
| | *nrf2 C7* | ACACAAACCACTGGAACCGT | TCTCCTCCAGGTCTGAGTCG |
| | *ncor1a* | GTCATCACCCCTCTCATGCCATC | CGTCGTCTCGATCGTGCTTG |
| | *ncor1b* | AGCACAAGCAGGACACACG | ACTGGTTCTCTGACCCTCCA |
| | *rab11Aab* | CGGGCCTTCGCAGAGAAAAAC | AGAATGGTCTGGAAAGCCGTT |
| | *rab11Ac* | CTTCCAGACCATTCTGACTGAAATC | GGTGACATGTTGTTGTCTTGCC |
| | *rab11Ad* | CTTCCAGACCATTCTGACTGAAATC | ACATGTCGTTGTCTCGTCG |
| | *rarαab* | AGCTCGGAAGAGATAGTCCCC | CTCCGGAAGAAACCCTTGCAG |

3 **Table S2.** List of up-regulated proteins in L2AC31-KO vs WT RTs

| Name of protein | Accession Number | log2(KO/WT) | log10(pval) |
|---|---|---|---|
| Isocitrate dehydrogenase (IDH) | A0A060W4K6 | 1.55 | 3.67 |
| motile sperm domain-containing protein 2 MOSPD2 | A0A060WQ66 | 1.27 | 3.65 |
| Isocitrate dehydrogenase (IDH) | A0A8C7QHT2 | 1.92 | 3.61 |
| Ubiquitin-conjugating enzyme E2 K (UBE2K) | A0A8C7V187 | 2.09 | 3.37 |
| Transferrin receptor 1b (Tr1b) | A0A060YMI7 | 1.31 | 3.35 |
| dhrs7b (Dehydrogenase/reductase 7) | A0A060WQL1 | 1.49 | 2.92 |
| SGT1 | A0A060YDC6 | 1.23 | 2.91 |
| very long-chain acyl-CoA synthetase ACSVL | A0A060YZL8 | 1.23 | 2.87 |
| Talin 1 | A0A060Y5X6 | 1.75 | 2.87 |
| RAD23A | A0A060YG13 | 4.05 | 2.80 |
| MFN2 (mitofusin 2) | A0A060YG13 | 3.81 | 2.77 |
| Glutamate–cysteine ligase (GCL) | A0A060WN13 | 1.44 | 2.51 |
| ZO-1 tight junction protein | A0A060YR10 | 1.04 | 2.47 |
| Prostaglandin E synthase 3 - chaperone p23 | A0A060XMQ8 | 1.14 | 2.41 |
| Galectin-8 LGALS8 | A0A060YPK1 | 1.34 | 2.26 |
| 28S ribosomal protein S35, mitochondrial (MRPS35) | A0A060XNY9 | 2.74 | 2.32 |
| TCB1 transposase | A0A060W525 | 1.35 | 2.26 |
| DP-N-acetylglucosamine pyrophosphorylase (UAP1) | A0A060WK17 | 1.00 | 2.11 |
| Aldose Reductase | A0A060W451 | 1.17 | 2.08 |
| Mitochondrial protein LACTB | A0A060WDJ0 | 1.34 | 1.94 |
| ALDOA - Aldolase - fructose-bisphosphate A | A0A060WUI1 | 1.18 | 1.90 |
| Saccharopine dehydrogenase (SDH) | A0A060YW21 | 1.28 | 1.83 |
| Phosphoserine transaminase - Phosphoserine Aminotransferase | A0A060YU89 | 1.51 | 1.82 |
| Mustn1 musculoskeletal embryonic nuclear protein 1a | A0A060XD24 | 2.08 | 1.82 |
| ALDOA - Aldolase - fructose-bisphosphate A | A0A060XHN5 | 1.16 | 1.80 |

| | | | |
|---|---|---|---|
| Condensin complex subunit 1 | A0A060XX58 | 1.33 | 1.78 |
| Vacuolar protein sorting-associated protein 33B - VPS33B | A0A060WA63 | 1.07 | 1.73 |
| Phosphoserine Aminotransferase | A0A060ZSW8 | 2.60 | 1.75 |
| tmem97 - Transmembrane Protein 97 | A0A060Y953 | 1.19 | 1.71 |
| MROH2B maestro heat like repeat family member 2B | A0A060XSP9 | 1.19 | 1.68 |
| Vacuolar protein sorting-associated protein 4B - VPS4B | A0A060X3F6 | 1.34 | 1.64 |
| ADP-ribosylation factor-like 3 (Arl3) | A0A060WFD0 | 1.59 | 1.64 |
| Nucleoporin 133 (Nup133) | A0A060X0R2 | 1.17 | 1.63 |
| mRNA-decapping enzyme 1A (DCP1A) | A0A060YNK5 | 1.26 | 1.62 |
| V-ATPase | A0A060XXM1 | 1.15 | 1.61 |
| Histone deacetylase (HDAC) | A0A060Z7V8 | 1.13 | 1.61 |
| Thioredoxin | C1BH85 | 3.08 | 1.59 |
| TRAPPC8 (Trafficking Protein Particle Complex Subunit 8) | A0A060WA90 | 3.77 | 1.58 |
| Elongation factor G 1, mitochondrial (GFM1) | A0A060VWB3 | 3.86 | 1.53 |
| PTPN23 (Protein Tyrosine Phosphatase Non-Receptor Type 23 | A0A060WXG6 | 1.23 | 1.53 |
| PRKAB1 - 5'-AMP-activated protein kinase subunit beta-1 | A0A060XTP5 | 1.49 | 1.51 |
| Influenza virus NS1A-binding protein (NS1BP) | A0A060W6C5 | 1.03 | 1.47 |
| Replication protein A (RPA) | A0A060XWW5 | 1.22 | 1.46 |
| ADP-ribosylhydrolase | A0A060XZF2 | 1.02 | 1.44 |
| Complement component 8 c8g | A0A060XV51 | 1.14 | 1.41 |
| 5-oxoprolinase | A0A060WE17 | 1.07 | 1.39 |
| Ubiquitin-protein ligase UBR5 | A0A060WTW8 | 2.63 | 1.42 |
| ATP citrate synthase (ACLY) | A0A060WTD0 | 1.69 | 1.38 |
| Glucose-6-phosphate 1-dehydrogenase (G6PD) | A0A060WHZ0 | 1.16 | 1.34 |
| Heat shock protein 75 kDa (TRAP1) | A0A060YT17 | 1.20 | 1.34 |
| FAM185A | A0A060WBH4 | 1.21 | 1.34 |
| 5-oxoprolinase | A0A060WDM6 | 1.38 | 1.30 |

| | | | |
|---|---|---|---|
| Hexokinase HK | A0A060WFB8 | 0.52 | 2.65 |
| G6PD (glucose-6-phosphate dehydrogenase | A0A060XEX6 | 0.77 | 1.57 |
| motile sperm domain-containing protein 2 | A0A674F7Q4 | 1.27 | 3.65 |
| Dual specificity phosphatase 23b (DUS23) | C1BIA0 | 0.86 | 3.59 |
| argininosuccinate synthase (ASS) | A0A060WCU7 | 0.52 | 3.39 |
| Integrator complex subunit 11 INTS11 | A0A060X0J0 | 0.55 | 1.66 |
| transferrin receptor 1b (Tr1b) | A0A060YPC2 | 1.31 | 3.35 |
| Trans-L-3-hydroxyproline dehydratase | A0A060WY07 | 0.85 | 3.10 |
| PTGES2 Gene - Prostaglandin E Synthase 2 | A0A060Z7I8 | 0.65 | 2.93 |
| Peroxisomal targeting signal 1 receptor (PTS1) | A0A060XU62 | 0.66 | 2.84 |
| ACTR1B Beta-centractin | D5LNF9 | 0.56 | 2.81 |
| Talin 1 | A0A060Y5X6 | 1.75 | 2.87 |
| RAD23 | A0A060Y182 | 3.81 | 2.77 |
| Malate dehydrogenase (MDH) | A0A8C7TVA0 | 0.55 | 2.57 |
| Malate dehydrogenase (MDH) | A0A060XXI1 | 0.55 | 2.54 |
| akr1a1a aldehyde reductase | A0A060W451 | 0.52 | 2.49 |
| Phosphoprotein enriched in astrocytes-15 (PEA-15) | A0A060Z5M9 | 0.85 | 2.47 |
| ZO-1 tight junction protein | A0A060YR10 | 1.04 | 2.47 |
| PTGES3 Gene - Prostaglandin E Synthase 3 | A0A060XMQ8 | 1.14 | 2.41 |
| Septin 9 (SEPT9) | A0A060X5R9 | 0.55 | 2.36 |
| Histone-lysine N-methyltransferase SETD5 | A0A8C7RCG9 | 0.86 | 2.33 |
| AFG3-like protein | A0A060X3F1 | 0.53 | 2.29 |
| Glutamate–cysteine ligase (GCL) | A0A8C7VYS9 | 1.35 | 2.26 |
| lgals8a galectin 8a | A0A8C7QLP1 | 1.34 | 2.26 |
| ATP6V1C1 V-type proton ATPase subunit C 1 | A0A060ZG28 | 0.54 | 2.21 |
| glrx glutaredoxin (thioltransferase) | A0A673ZI75 | 0.58 | 2.21 |
| 3alpha(or 20beta)-hydroxysteroid dehydrogenase | A0A674DT84 | 0.77 | 2.16 |

| | | | |
|---|---|---|---|
| ABCB6 ATP-binding cassette super-family B member 6, mitochondrial | A0A060X8Q2 | 0.98 | 1.35 |
| Influenza virus NS1A-binding protein (NS1BP) | A0A060WEI1 | 0.67 | 1.31 |
| Synaptobrevin homolog YKT6 | A0A060XS36 | 0.58 | 1.32 |
| unknown | A0A060WAB1 | 0.60 | 1.32 |
| Mitochondrial amidoxime-reducing component 1 MOSC1 | A0A060Z7S4 | 0.63 | 1.33 |
| Mitochondrial outer-membrane protein FUNDC1 | A0A060WG90 | 0.67 | 1.34 |
| GSTO1 (Glutathione S-transferase omega) GSTTL | C1BFM5 | 0.73 | 1.39 |
| Transaldolase | A0A060YV08 | 0.58 | 1.34 |
| Protein ETHE1, mitochondrial | A0A8C7SHX1 | 0.92 | 1.41 |
| Exocyst complex component 1 EXOC1 | A0A060X3B4 | 0.93 | 1.40 |
| mao Amine oxidase | P49253 | 0.73 | 1.43 |
| Exocyst complex component 1 EXOC1 | A0A060X3Y7 | 0.78 | 1.45 |
| ARHGAP1 (Rho GTPase Activating Protein 1 | A0A8U0QMV8 | 0.81 | 1.46 |
| NMRK1 nicotinamide riboside kinase 1 | A0A060WPZ9 | 0.58 | 1.39 |
| Protein transport protein Sec24B | A0A060W285 | 0.54 | 1.38 |
| Protein kinase C PKC | A0A060X6F9 | 0.51 | 1.39 |
| Kinesin-1 KIF5B | A0A060WML3 | 0.60 | 1.41 |
| Thioredoxin reductases (TR | A0A060XUV9 | 0.62 | 1.42 |
| Methylmalonyl CoA epimerase MCEE | A0A060XNR5 | 0.56 | 1.45 |
| Palmitoyl-protein hydrolase (lyplal1) | A0A060YP76 | 0.61 | 1.44 |
| Probable ubiquitin carboxyl-terminal hydrolase FAF-X | A0A060XT85 | 0.64 | 1.45 |
| 2',3'-cyclic-nucleotide 3'-phosphodiesterase CNPase | A0A060W1Z9 | 0.65 | 1.47 |
| Phosphatidylinositol transfer protein beta isoform PITPNB | A0A060XQV7 | 0.70 | 1.47 |
| nudC domain-containing protein 3 | A0A060Y4N3 | 0.76 | 1.50 |
| atp6v1f ATPase H+ transporting V1 subunit F | A0A8C7QNX0 | 0.66 | 1.50 |
| PTGES2 Gene - Prostaglandin E Synthase 2 | A0A060YE60 | 0.50 | 1.51 |
| Fibronectin FN | A0A060YRG9 | 0.54 | 1.55 |

| | | | |
|---|---|---|---|
| Fibulin-1 FBLN1 | A0A060XB63 | 0.72 | 1.53 |
| GLRX5 Glutaredoxin 5 homolog | C1BI21 | 0.72 | 1.55 |
| coenzyme Q : cytochrome c – oxidoreductase COMPLEX III | A0A060Z810 | 0.81 | 1.55 |
| cAMP-regulated phosphoprotein 21 ARPP-21 | A0A060Y8E9 | 0.61 | 1.58 |
| PATL1 | A0A060WEQ7 | 0.51 | 1.59 |
| Calponin 2 | A0A060YZQ0 | 0.78 | 1.60 |
| bri3bp | A0A060XE57 | 0.59 | 1.62 |
| cysS cysteine--tRNA ligase | A0A060W354 | 0.66 | 1.63 |
| SLC27A4 | A0A060XKK5 | 0.90 | 1.62 |
| Histone deacetylase HDAC | A0A060Z7V8 | 1.13 | 1.61 |
| V-ATPase | A0A060XXM1 | 1.15 | 1.61 |
| 5'-(N(7)-methylguanosine 5'-triphospho)-[mRNA] hydrolase / D10 decapping enzyme | A0A060YNK5 | 1.26 | 1.62 |
| Heat Shock Protein Family A (Hsp70) Member 14 HSPA14 | A0A060Y0Z6 | 0.57 | 1.67 |
| Threo-3-hydroxyaspartate ammonia-lyase D-THA DH | A0A060YKP0 | 0.63 | 1.67 |
| rcor1 REST corepressor 1 | A0A674AG29 | 0.86 | 1.68 |
| Isocitrate dehydrogenase IDH | A0A060Z6S7 | 0.92 | 1.69 |
| Vacuolar protein sorting-associated protein 33B - VPS33B | A0A060WA63 | 1.07 | 1.73 |
| Proteoglycan 4 or lubricin | A0A060Z3V5 | 0.89 | 1.80 |
| aifm2 Apoptosis inducing factor mitochondria associated 2 | A0A060W214 | 0.53 | 1.72 |
| Presequence protease (PreP | A0A060WBN1 | 0.52 | 1.76 |
| Dihydrolipoyl transacetylase dlat-1 | A0A061A6R7 | 0.59 | 1.77 |
| E3 ubiquitin-protein ligase NEDD4 | A0A060Y4F6 | 0.73 | 1.81 |
| Secretory carrier-associated membrane protein 2 (SCAMP2) | A0A060W547 | 0.72 | 1.82 |
| Dynein cytoplasmic 1 heavy chain 1 DYNC1H1 | A0A060XFE1 | 0.58 | 1.84 |
| 26S protease regulatory subunit 4 PSMC1 | A0A060XW34 | 0.51 | 1.84 |
| LIMA1 LIM domain and actin-binding protein 1 | A0A060YJ77 | 0.72 | 1.89 |

| | | | |
|---|---|---|---|
| RNA helicase MOV-10 | A0A8C7LVX8 | 0.52 | 1.91 |
| Caspase 3, apoptosis-related cysteine peptidase b | A0A060W2H2 | 0.88 | 1.96 |
| TIP41-like protein | A0A060X165 | 0.68 | 2.01 |
| Cdc42 effector protein 4 CDC42EP4 | A0A060Z1R2 | 0.62 | 2.01 |
| fatty-acid-binding protein (FABP) | A0A060YTS7 | 1.00 | 2.03 |
| usp11 Ubiquitinyl hydrolase 1 | A0A060XEN3 | 0.71 | 2.06 |
| ACOT9 Acyl-CoA thioesterase 9 | A0A060Y438 | 0.62 | 2.07 |
| TIP41-like protein | A0A060WZQ8 | 0.83 | 2.05 |
| UDP-N-acetylhexosamine pyrophosphorylase UAP1 | A0A8C7UHI6 | 0.97 | 2.07 |
| C-factor | A0A060WU91 | 0.69 | 2.10 |
| UDP-N-acetylhexosamine pyrophosphorylase UAP1 | A0A060WK17 | 1.00 | 2.11 |
| Aldo-keto reductase family 1, member B1 (AKR1B1) - Aldose reductase | A0A060W451 | 1.17 | 2.08 |

**Table S3.** List of down-regulated proteins in L2AC31-KO vs WT RTs

| Name of protein | Accession | log2(KO/WT) | log10(pval) |
|---|---|---|---|
| apolipoproteinB-100 | A0A060W3H4 | -1.61 | 4.62 |
| apolipoproteinB-100 | A0A060Y552 | -3.23 | 4.12 |
| homogentisate 1,2-dioxygenase (HGD) | A0A060YA42 | -1.18 | 3.75 |
| Carbonic anhydrase 1 (CA) | Q68YC2 | -1.29 | 3.49 |
| apolipoproteinB-100 | A0A060Z709 | -3.21 | 2.92 |
| Sodium-coupled neutral amino acid transporter 4 (SNAT4) | A0A060X0J3 | -1.17 | 2.85 |
| Surfeit locus protein 2 SURF2 | A0A060W7Q0 | -1.61 | 2.79 |
| D-aspartate oxidase | A0A060WJY0 | -1.80 | 2.59 |
| Gastrula zinc finger protein | A0A140F304 | -1.64 | 2.56 |
| venom factor isoform X1 | A0A060XW02 | -2.06 | 2.46 |
| tRNA methyltransferase | A0A060WFW7 | -2.61 | 2.38 |
| Retinol dehydrogenase 12 | A0A060X190 | -2.15 | 2.34 |
| SELENBP1 / methanethiol oxidase | A0A060XQH4 | -1.18 | 2.38 |
| ZNT7 Zinc transporter protein 7 | A5PMX1 | -1.05 | 2.32 |
| Vitellogenin 1 | Q92093 | -2.02 | 2.19 |
| Retinol-binding protein 2 (RBP2) | Q71B03 | -1.09 | 2.17 |
| MHC class I | A0A060WZV0 | -1.43 | 2.13 |
| Terminal uridylyl transferase 7 (TUT7) | A0A060YJN8 | -1.09 | 2.13 |
| Tropomodulin 3 | A0A060ZI14 | -1.91 | 2.10 |
| Immunoglobulin kappa light chain | A0A060XIK2 | -1.48 | 2.08 |
| Sulfide:quinone oxidoreductase (SQOR) | A0A060YB67 | -1.50 | 2.06 |
| GSTM3 glutathione S-transferase mu 3 | A0A060Y2H6 | -1.23 | 2.03 |
| C1-inhibitor (C1-inh) | A0A060WER0 | -1.47 | 1.99 |
| Ribosome-binding protein 1 (p180) | A0A060VRW0 | -1.02 | 1.96 |
| GTPase IMAP family member 8 (GIMA8) | A0A060YJA9 | -1.32 | 1.94 |
| Monocarboxylate transporter 7 (MCT7) | A0A060XZE8 | -1.55 | 1.93 |

| | | | |
|---|---|---|---|
| inactive rhomboid protein 2 (iRhom2) | A0A060WAR8 | -1.46 | 1.91 |
| Cu transport protein Antioxidant-1 (Atox1) | A0A060YR00 | -1.01 | 1.90 |
| NPC2 NPC intracellular cholesterol transporter 2 | C1BGA1 | -1.03 | 1.85 |
| wdr92 - WD repeat-containing protein 92 | A0A060XN54 | -1.07 | 1.84 |
| Phosphorylase kinase (PhK) | A0A060VME0 | -1.05 | 1.83 |
| Cathepsin F | A0A060WE48 | -1.07 | 1.83 |
| Aromatic L-amino acid decarboxylase (AADC or AAAD) DOPA decarboxylase (DDC) | A0A060WI79 | -1.53 | 1.80 |
| Carboxypeptidase D | A0A060XHB4 | -1.27 | 1.76 |
| N-acetyltransferase 10 - NAT-10 | A0A060WTM3 | -1.55 | 1.75 |
| PLRG1 - Pleiotropic regulator 1 | A0A060XMA6 | -1.31 | 1.73 |
| Acyl-CoA thioesterase - ACOT | A0A060XFI2 | -1.44 | 1.71 |
| Guanylate-binding protein 1 (GBP1) | A0A060X412 | -2.74 | 1.69 |
| dnttip2 deoxynucleotidyltransferase | A0A060WM03 | -1.96 | 1.68 |
| Butyrophilin subfamily 1 member A1 (BTN1A1) | A0A060X431 | -1.36 | 1.67 |
| Properdin | A0A060XDG4 | -1.59 | 1.66 |
| ZCHC4 - rRNA N6-methyltransferase | A0A060Y1B0 | -1.03 | 1.63 |
| FAD-dependent oxidoreductase 1 (FOXRED1) | A0A060VXJ0 | -1.29 | 1.62 |
| HIUHase, and 5-hydroxyisourate hydrolase | A0A8C7TKN0 | -1.27 | 1.61 |
| Golgi membrane protein 1 (GOLM1) | A0A060WNZ1 | -1.01 | 1.38 |
| Nucleobindin-2 - NUCB2 | A0A060WJ28 | -1.08 | 1.60 |
| SLC27A6 - Long-chain fatty acid transport protein 6 | A0A060YFS1 | -2.15 | 1.60 |
| Cytochrome c oxidase polypeptide VIIc | P80334 | -1.17 | 1.59 |
| PHYIP Phytanoyl-CoA hydroxylase-interacting protein | A0A060XE45 | -2.29 | 1.54 |
| Prolyl isomerase (also known as peptidylprolyl isomerase or PPIase) | A0A060VQQ6 | -1.28 | 1.52 |
| Synaptosomal-Associated Protein, 25kDa (SNAP-25) | A0A060WCC0 | -1.64 | 1.48 |
| Cell adhesion molecule 1 (CADM1) | A0A060VYC8 | -1.36 | 1.48 |
| ubiquitin-like protein FUBI | C1BFG2 | -1.31 | 1.48 |

| | | | |
|---|---|---|---|
| CDGSH iron sulfur domain | C1BI29 | -1.16 | 1.42 |
| COMM domain-containing protein 9 | A0A060XCH2 | -1.34 | 1.35 |
| GSTM3 - Glutathione S-transferase Mu 3 | A0A060ZAG6 | -1.00 | 1.34 |
| NADH dehydrogenase (ubiquinone) 1 alpha NDUFA7 | A0A060W5N8 | -1.35 | 1.34 |
| ACIN1 (Apoptotic Chromatin Condensation Inducer 1) | A0A060XGP3 | -1.16 | 1.33 |
| GPRC5C G-protein coupled receptor family C group 5 member C | A0A060XR44 | -1.45 | 1.32 |
| ACIN1 (Apoptotic Chromatin Condensation Inducer 1) | A0A060Y2W1 | -1.36 | 1.33 |
| EARP-interacting protein homolog | A0A060XAA8 | -1.01 | 1.31 |
| Y-box binding protein YB-1 | A0A060Y0T3 | -0.77 | 1.31 |
| UDP-glucuronosyltransferase 2A3 UGT2A3 | A0A060YBS5 | -0.90 | 1.32 |
| EARP-interacting protein homolog | A0A060XAA8 | -1.01 | 1.31 |
| Cell division cycle and apoptosis regulator protein 1 CCAR1 | A0A060XZF0 | -0.98 | 1.32 |
| Calcium/calmodulin-dependent protein kinase type 1 CAMK1 | A0A060YCX0 | -0.61 | 1.35 |
| Thiopurine methyltransferase TPMT | A0A060WUW1 | -0.73 | 1.37 |
| Activating signal cointegrator 1 (ASC-1) | A0A060Z7U5 | -0.68 | 1.40 |
| ctsh cathepsin H | A0A8C7S3U7 | -0.62 | 1.40 |
| scavenger receptor class B type I (SR-BI) | A0A060WH67 | -0.57 | 1.39 |
| Sarcosine dehydrogenase | A0A060YZQ1 | -0.58 | 1.41 |
| OS9 endoplasmic reticulum lectin ERLEC1 | A0A8C7TGM4 | -0.53 | 1.40 |
| Protein kinase C and casein kinase substrate PACSIN3 | A0A060VSI0 | -0.51 | 1.40 |
| GTP-binding nuclear protein Ran | A0A060VZL7 | -0.89 | 1.37 |
| Carboxypeptidase Q | A0A060YCQ8 | -0.89 | 1.41 |
| UDP-glucuronosyltransferase | A0A060YHW2 | -0.93 | 1.41 |
| Estradiol 17-beta-dehydrogenase 8 HSD17B8 | A0A060VRD9 | -0.83 | 1.43 |
| NADH dehydrogenase [ubiquinone] 1 beta NDUFB6 | A0A060XUW9 | -0.83 | 1.44 |
| BTD Biotinidase | A0A060XNN4 | -0.74 | 1.47 |

| | | | |
|---|---|---|---|
| Lactoylglutathione lyase | A0A060WQQ3 | -0.71 | 1.48 |
| NDUFB11 NADH dehydrogenase 1 beta subcomplex subunit 11 | A0A060XEM0 | -0.56 | 1.47 |
| Glycoprotein endo-alpha-1,2-mannosidase MANEA | A0A060XJZ5 | -0.58 | 1.48 |
| CD276 antigen-like | A0A060WBI1 | -0.66 | 1.48 |
| vtg1 Vitellogenin | Q92093 | -2.02 | 2.19 |
| Sulfide:quinone oxidoreductase (SQOR) | A0A060ZDL1 | -0.91 | 2.78 |
| Organic solute transporter subunit alpha (SLC51A) or OST-alpha | A0A060XRH2 | -0.75 | 2.82 |
| pm20d1 N-fatty-acyl-amino acid synthase/hydrolase | A0A060XQK1 | -0.87 | 2.64 |
| Ras-like GTP-binding protein Rho1 | A0A060X4I6 | -0.57 | 3.63 |
| Lysosomal-associated membrane protein 1 (LAMP-1) | A0A8C7VQL2 | -1.64 | 2.56 |
| TM9SF2 transmembrane 9 superfamily member 2 | A0A674CL93 | -0.60 | 2.37 |
| ndufv1 NADH:ubiquinone oxidoreductase | A0A674CHH5 | -0.61 | 2.34 |
| cytosolic non-specific dipeptidase CNDP2 | B5X4T7 | -0.54 | 2.28 |
| Prefoldin subunit 5 PFDN5 | A0A060WP50 | -0.50 | 2.25 |
| GDP-L-fucose synthase | A0A060YFS3 | -0.66 | 2.24 |
| POLR1C DNA-directed RNA polymerases I and III subunit RPAC1 | A0A060WQ06 | -0.77 | 2.24 |
| Monocarboxylate transporter 10 SLC16A10 | A0A060WBK6 | -0.74 | 2.20 |
| Integrin beta-1 (ITGB1), also known as CD29 | A0A060X309 | -0.65 | 2.18 |
| Membrane-associated progesterone receptor component 2 PGRMC2 | A0A8U0Q940 | -0.61 | 2.17 |
| Sarcosine dehydrogenase | A0A060YK79 | -0.51 | 2.17 |
| PRMT1 protein arginine methyltransferase 1 | A0A060X602 | -0.55 | 2.14 |
| DHRS4 - Dehydrogenase/reductase SDR family member 4 | A0A8U0QWT5 | -0.63 | 2.13 |
| Adenylate cyclase | A0A060WN48 | -0.58 | 2.08 |
| TPK2 - Thiamine pyrophosphokinase 2 | A0A060Z2Y2 | -0.51 | 2.08 |
| PTER - Phosphotriesterase-related protein | A0A060Y8N1 | -0.62 | 2.02 |
| 60S acidic ribosomal protein P2 | A0A060WJW3 | -0.67 | 2.02 |

| | | | |
|---|---|---|---|
| DPYD dihydropyrimidine dehydrogenase | A0A060YFT0 | -0.79 | 2.02 |
| ACY3 Aspartoacylase | A0A060WF45 | -0.94 | 1.99 |
| CRYL1 Crystallin, lambda 1 or L-gulonate 3-dehydrogenase | A0A060XAB5 | -0.80 | 1.97 |
| GTP cyclohydrolase I feedback regulatory protein GFRP | C1BHD1 | -0.69 | 1.96 |
| GSTK1 glutathione S-transferase kappa 1 | A0A060XKR8 | -0.59 | 1.98 |
| phr2aB Serine/threonine-protein phosphatase 2A regulatory subunit B (PP2C) | A0A060XGH8 | -0.58 | 1.97 |
| PXMP2 - Peroxisomal membrane protein 2 | A0A060WGG2 | -0.87 | 1.92 |
| TM9SF2 transmembrane 9 superfamily member 2 | A0A8K9V045 | -0.55 | 1.94 |
| FAHD2A (Fumarylacetoacetate Hydrolase Domain Containing 2A) | A0A060W664 | -0.74 | 1.85 |
| Malate synthase | A0A060XNA5 | -0.52 | 1.85 |
| Hepatocyte nuclear factor 4 alpha (HNF4A) | A0A060X563 | -0.77 | 1.83 |
| Indoleamine 2,3-dioxygenase 2 (IDO2) | A0A060VVQ1 | -0.96 | 1.81 |
| KH domain-containing, RNA-binding, signal transduction-associated protein 1 (KHDRBS1) | A0A060YV02 | -0.69 | 1.74 |
| cysteine-rich with EGF-like domains 2 (CRELD2) | A0A8C7VXU9 | -0.84 | 1.75 |
| MYEF2 myelin expression factor 2 | A0A061A2A1 | -0.96 | 1.75 |
| Acyl-CoA thioesterase 16 ACOT | A0A060YTU9 | -0.89 | 1.72 |
| Peroxidase | A0A060XUD1 | -0.62 | 1.70 |
| Atp11c - Phospholipid-transporting ATPase 11C | A0A060XC13 | -0.52 | 1.69 |
| Alpha-1-microglobulin/bikunin precursor AMBP | A0A060YFZ1 | -0.63 | 1.68 |
| Aldolase B ALDOB | A7L6A8 | -0.62 | 1.67 |
| Transportin-1 | A0A060Y475 | -0.57 | 1.65 |
| Integrin beta-1 (ITGB1) | A0A060WSX3 | -0.88 | 1.68 |
| UDP-glucuronosyltransferase 2A1 UGT2A1 | A0A060Y8W2 | -0.94 | 1.65 |
| Cytochrome c oxidase polypeptide VIIc cox-7c | P80334 | -1.17 | 1.59 |
| Nucleobindin-2 - NUCB2 | A0A060WJ28 | -1.08 | 1.60 |

| | | | |
|---|---|---|---|
| Dihydropyrimidinase | A0A060ZNP9 | -0.93 | 1.62 |
| Desmoplakin a | A0A060VWG6 | -0.82 | 1.62 |
| Glycerol-3-phosphate dehydrogenase (GPDH) | A0A8C7P1A0 | -0.61 | 1.63 |
| UDP-glucose 6-dehydrogenase | A0A060WVE9 | -0.95 | 1.56 |
| uncharacterized protein LOC110526684 | A0A060WA68 | -0.78 | 1.55 |
| Low-density lipoprotein receptor-related protein-1 (LRP1) | A0A060XD42 | -0.71 | 1.54 |
| Pyridoxine 5'-phosphate synthase DXP | A0A060W490 | -0.60 | 1.53 |
| cysteine-rich with EGF-like domains 2 (CRELD2) | A0A8C7VXU9 | -0.71 | 1.61 |
| UFM1 ubiquitin fold modifier 1 | A0A060YYP8 | -0.60 | 1.61 |
| NOL5A Nucleolar protein 56 | A0A060W4J3 | -0.67 | 1.60 |
| U6 snRNA-associated Sm-like protein LSm7 | C1BH51 | -0.64 | 1.57 |
| Aldo-keto reductase family 1, member B1 (AKR1B1) - Aldose reductase | A0A060ZCT9 | -0.61 | 1.56 |

**Table S4.** List of differentially-regulated proteins in L2AC31-KO vs WT RTs mentioned and potential KFERQ-like motifs

Upregulated proteins

| Name of protein | Accession number | log2(KO/WT) | log10(pval) | KFERQ-like motif |
|---|---|---|---|---|
| Isocitrate dehydrogenase (IDH) | A0A060W4K6 | 1.55 | 3.67 | 1 canon. 1 acet. |
| Isocitrate dehydrogenase (IDH) | A0A8C7QHT2 | 1.92 | 3.61 | 1 acet. 1 phos. |
| Isocitrate dehydrogenase (IDH) | A0A060Z6S7 | 0.92 | 1.69 | 1 acet. |
| Ubiquitin-conjugating enzyme E2 K (UBE2K) | A0A8C7V187 | 2.09 | 3.37 | 3 canon. 4 acet. |
| very long-chain acyl-CoA synthetase ACSVL | A0A060YZL8 | 1.23 | 2.87 | 1 canon. |
| RAD23 homolog A | A0A060Y182 | 3.81 | 2.77 | 4 phos. 1 acet. |
| RAD23 homolog A | A0A060YG13 | 4.05 | 2.80 | 4 phos. 1 acet. |
| Aldolase - fructose-bisphosphate (ALDOA) | A0A060WUI1 | 1.18 | 1.90 | 1 canon. 2 phos. 1 acet. |
| Aldolase - fructose-bisphosphate A (ALDOA) | A0A060XHN5 | 1.16 | 1.80 | 1 canon. 1 phos. 1 acet. |
| Vacuolar protein sorting-associated protein 33B (VPS33B) | A0A060WA63 | 1.07 | 1.73 | 1 canon. |
| Vacuolar protein sorting-associated protein 4B (VPS4B) | A0A060X3F6 | 1.34 | 1.64 | 1 phos. 4 acet. |
| Thioredoxin | C1BH85 | 3.08 | 1.59 | none |
| Protein Tyrosine Phosphatase Non-Receptor Type 23 (PTPN23) | A0A060WXG6 | 1.23 | 1.53 | 2 canon. 6 phos. 2 acet. |
| Thioredoxin reductases (TR) | A0A060XUV9 | 0.62 | 1.42 | 1 phos. 2 acet. |
| ATP citrate synthase (ACLY) | A0A060WTD0 | 1.69 | 1.38 | none |
| Glucose-6-phosphate 1-dehydrogenase (G6PD) | A0A060WHZ0 | 1.16 | 1.34 | none |
| Glucose-6-phosphate 1-dehydrogenase (G6PD) | A0A060XEX6 | 0.77 | 1.57 | 1 canon. 1 phos. 1 acet. |
| Hexokinase (HK) | A0A060WFB8 | 0.52 | 2.65 | 5 canon. 6 phos. 1 acet. |

| | | | | |
|---|---|---|---|---|
| Malate dehydrogenase (MDH) | A0A8C7TVA0 | 0.55 | 2.57 | none |
| Malate dehydrogenase (MDH) | A0A060XXI1 | 0.55 | 2.54 | none |
| Glutamate–cysteine ligase (GCL) | A0A8C7VYS9 | 1.35 | 2.26 | 1 canon. |
| phosphofructokinase-2 (PFK-2) | A0A060Y9T0 | 0.51 | 1.30 | 1 canon. |
| Vacuolar protein sorting-associated protein 33B (VPS33B) | A0A060WA63 | 1.07 | 1.73 | 1 canon. |
| 26S protease regulatory subunit 4 (PSMC1) | A0A060XW34 | 0.51 | 1.84 | 1 canon. 1 acet. |
| fatty-acid-binding protein (FABP) | A0A060YTS7 | 1.00 | 2.03 | none |

Downregulated proteins

| Name of protein | Accession number | log2(KO/WT) | log10(pval) | KFERQ-like motif |
|---|---|---|---|---|
| apolipoproteinB-100 | A0A060W3H4 | -1.61 | 4.62 | 9 canon. 3 phos. 8 acet. |
| apolipoproteinB-100 | A0A060Y552 | -3.23 | 4.12 | none |
| apolipoproteinB-100 | A0A060Z709 | -3.21 | 2.92 | 3 phos. 1 acet. |
| FAD-dependent oxidoreductase 1 (FOXRED1) | A0A060VXJ0 | -1.29 | 1.62 | 1 canon. 1 acet. |
| Adenylate cyclase | A0A060WN48 | -0.58 | 2.08 | 4 canon. 3 phos. 1 acet. |

**Table S5.** High Carbohydrates diet composition

| Ingredients % | Diet<br>**HighCHO** | |
|---|---|---|
| Fish meal | 60.6 | ***1*** |
| Starch | 30 | ***2*** |
| Fish oil | 5 | ***3*** |
| Alginate | 2 | ***4*** |
| Vitamin mix | 1.2 | ***5*** |
| Mineral mix | 1.2 | ***6*** |
| Proximate composition | 95.97 | |
| Dry matter (DM) %diet | 43.9 | |
| Crude protein %DM | 11.47 | |
| Crude lipid %DM | 24.66 | |
| Gross energy %DM | 20.96 | |
| Ash %DM | 12.83 | |

**Table S6.** List of other primers used in study (forward and reverse)

| Name of gene | Forward sequence | Reverse sequence |
|---|---|---|
| *L2b C14* | GAATTTGCTACAGCCCATGA | GTCATTAATGGAAAGATTACACAGC |
| *L2b C31* | GAATTTGCTACAGCCCATGA | CACCCCACACAAGGATCAAC |
| *L2c C14* | ATTTGCTACAGCCGTGGAGT | GGAGCCAACGTGTTGTGAC |
| *L2c C31* | ATTTGCTACAGCCGTGGAGT | GCAACAGGGAACATGTTGTG |
| *L1 C3* | GCAGCAGAGAGAGCACAAG | CTTGCGGCTATGTTGTTGTG |
| *L1 C22* | TTGTGTGCATGCTGTTGTGA | CCATTGCCGTTGGTTACAGT |
| *GCK-a* | CTGCCCACCTACGTCTGT | GTCATGGCGTCCTCAGAGAT |
| *GCK-b* | TCTGTGCTAGAGACAGCCC | CATTTTGACGCTGGACTCCT |
| *PFK* | GATCCCTGCCACCATCAGTA | GTAACCACAGTAGCCTCCCA |
| *GAPDH* | CTGGAGAAGCCTGCCAGCTA | CCATGTGACCAGCTTGACGA |
| *PK* | CCATCGTCGCGGTAACAAGA | GCCCCTGGCCTTTCCTATGT |
| *Me1* | TACGTGCGGTGTGTGTGACG | GTGCCCACATCCAGCATGAC |
| *ACLY* | GCTTTTGCCACGGTGGTCTC | GCTTCCGCTACGCCAATGTC |
| *G6PD* | CTCATGGTCCTCAGGTTTG | AGAGAGCATCTGGAGCAAGT |
| *FASN* | TGATCTGAAGGCCCGTGTCA | GGGTGACGTTGCCGTGGTAT |
| *GSR* | CTAAGCGCAGCGTCATAGTG | ACACCCCTGTCTGACGACAT |
